# Post-Transcriptional Control of Coenzyme Q Biosynthesis Revealed by Transomic Analysis of the RNA-Binding Protein Puf3p

**DOI:** 10.1101/146985

**Authors:** Christopher P. Lapointe, Jonathan A. Stefely, Adam Jochem, Paul D. Hutchins, Gary M. Wilson, Nicholas W. Kwiecien, Joshua J. Coon, Marvin Wickens, David J. Pagliarini

**Affiliations:** Department of Biochemistry, University of Wisconsin–Madison, Madison, WI 53706, USA; Morgridge Institute for Research, Madison, WI 53715, USA; Genome Center of Wisconsin, Madison, WI 53706, USA; Department of Chemistry, University of Wisconsin–Madison, Madison, WI 53706, USA; Department of Biomolecular Chemistry, University of Wisconsin–Madison, Madison, WI 53706, USA

## Abstract

Coenzyme Q (CoQ) is a redox active lipid required for mitochondrial oxidative phosphorylation (OxPhos). How CoQ biosynthesis is coordinated with the biogenesis of OxPhos protein complexes is unclear. Here, we show that the *Saccharomyces cerevisiae* RNA-binding protein (RBP) Puf3p directly regulates CoQ biosynthesis. To establish the mechanism for this regulation, we employed a transomic strategy to identify mRNAs that not only bind Puf3p, but also are regulated by Puf3p *in vivo*. The CoQ biosynthesis enzyme Coq5p is a critical Put3p target: Puf3p regulates the level of Coq5p and prevents its toxicity, thereby enabling efficient CoQ production. In parallel, Puf3p represses a specific set of proteins involved in mitochondrial protein import, translation, and OxPhos complex assembly — pathways essential to prime mitochondrial biogenesis. Our data reveal a mechanism for post-transcriptionally coordinating CoQ production with OxPhos biogenesis and, more broadly, demonstrate the power of transomics for defining genuine targets of RBPs.

**HIGHLIGHTS:**

- The RNA binding protein (RBP) Puf3p regulates coenzyme Q (CoQ) biosynthesis
- Transomic analysis of RNAs, proteins, lipids, and metabolites defines RBP targets
- Puf3p regulates the potentially toxic CoQ biosynthesis enzyme Coq5p
- Puf3p couples regulation of CoQ with a broader program for controlling mitochondria

## INTRODUCTION

Mitochondria are complex organelles central to cellular metabolism, and their dysfunction is implicated in over 150 human diseases (Vafai and Mootha, 2012). Alterations in metabolic demands induce adaptive changes in mitochondrial mass and composition (Labbe et al., 2014), but precisely how these changes are regulated is unclear. Transcriptional regulators of mitochondria have been identified (Kelly and Scarpulla, 2004), yet the roles of post-transcriptional regulators remain poorly defined.

Remodeling and biogenesis of mitochondria is uniquely complicated. It requires synchronized expression of proteins encoded by both nuclear DNA and mitochondrial DNA (mtDNA) (Couvillion et al., 2016), and coordinated assembly of protein complexes, such as those of oxidative phosphorylation (OxPhos), with specific metabolites and lipids, such as coenzyme Q (CoQ). CoQ is a redox active lipid in the electron transport chain whose deficiency causes numerous human diseases (Laredj et al., 2014). Its biosynthesis likewise requires assembly of a multi-protein complex of enzymes (“complex Q”) and lipids in the mitochondrial matrix (Floyd et al., 2016; He et al., 2014; Stefely et al., 2016b), and production of the water-soluble CoQ head group precursor 4-hydroxybenzoate (4-HB) (Payet et al., 2016; Stefely et al., 2016a). Thus, mitochondrial biogenesis demands integrated remodeling of the proteome, the metabolome, and the lipidome. The mechanisms for this “transomic” regulation are largely uncharacterized.

As described in this report, analysis of our publicly available “Y3K” data set (Stefely et al., 2016a) reveals an unexpected link between CoQ abundance and the *Saccharomyces cerevisiae* gene *puf3*. The encoded protein, Puf3p, belongs to the conserved PUF family of RNA-binding proteins (RBPs), which play key roles throughout Eukarya (Quenault et al., 2011; Spassov and Jurecic, 2003; Wickens et al., 2002). Puf3p binds to mRNAs containing “Puf3p-binding elements” (PBEs), conforming to the consensus UGUANAUA (Gerber et al., 2004; Olivas and Parker, 2000; Zhu et al., 2009), and regulates their translation through a variety of mechanisms (Houshmandi and Olivas, 2005; Jackson et al., 2004; Lee and Tu, 2015; Rowe et al., 2014). mRNAs that bind Puf3p have been identified, but which of those mRNAs are genuine “targets” – mRNAs whose activity *in vivo* is controlled by Puf3p – is an open question. The few established targets have connections to OxPhos complexes (e.g., *cox17*) and the mitochondrial ribosome (Chatenay-Lapointe and Shadel, 2011; Garcia-Rodriguez et al., 2007; Olivas and Parker, 2000). Accordingly, yeast that lack *puf3* (“Δ*puf3* yeast”) exhibit mitochondria-related phenotypes, including reduced respiratory growth (Eliyahu et al., 2010; Gerber et al., 2004; Lee and Tu, 2015) and increased respiratory activity during fermentation (Chatenay-Lapointe and Shadel, 2011). However, it is unclear whether Puf3p affects CoQ directly, via control of a key CoQ biosynthesis pathway enzyme, or indirectly, by controlling a protein that subsequently affects the pathway.

Here, we employ a transomic strategy that analyzes mRNAs, proteins, lipids, and metabolites to define high-confidence Puf3p targets, and we demonstrate that Puf3p directly regulates CoQ biosynthesis. In addition to defining a specific Puf3p target responsible for CoQ control—Coq5p—we reveal 90 additional high-confidence direct targets of Puf3p, which provides insight into how CoQ production is coordinated with additional OxPhos biogenesis pathways. The results also demonstrate the power of transomic strategies for dissecting the molecular functions of RBPs, which have widespread roles in human health and disease (Gerstberger et al., 2014).

## RESULTS

### Puf3p Regulates CoQ Biosynthesis

The Y3K data set (Stefely et al., 2016a) reveals Δ*puf3* yeast to be significantly (p < 0.05) deficient for CoQ when cultured under fermentation conditions, but not under respiration conditions (Figures 1A, 1B, S1A, and S1B). Notably, the early CoQ biosynthesis intermediate polyprenylhydroxybenzoate (PPHB) was elevated, while the later intermediate demethoxy-CoQ (DMQ) was decreased (Figure 1A), suggesting a defect in one of the steps in the CoQ pathway that depends on the CoQ biosynthetic complex (“complex Q” or “CoQ-Synthome”; comprised of Coq3p–Coq9p) (Figure 1B). The accumulation of PPHB in Δ*puf3* yeast was the largest such change across all yeast strains in the Y3K study under fermentation growth conditions (Figure S1A), further suggesting a functionally important link between Puf3p and the CoQ pathway.

**Figure 1.**
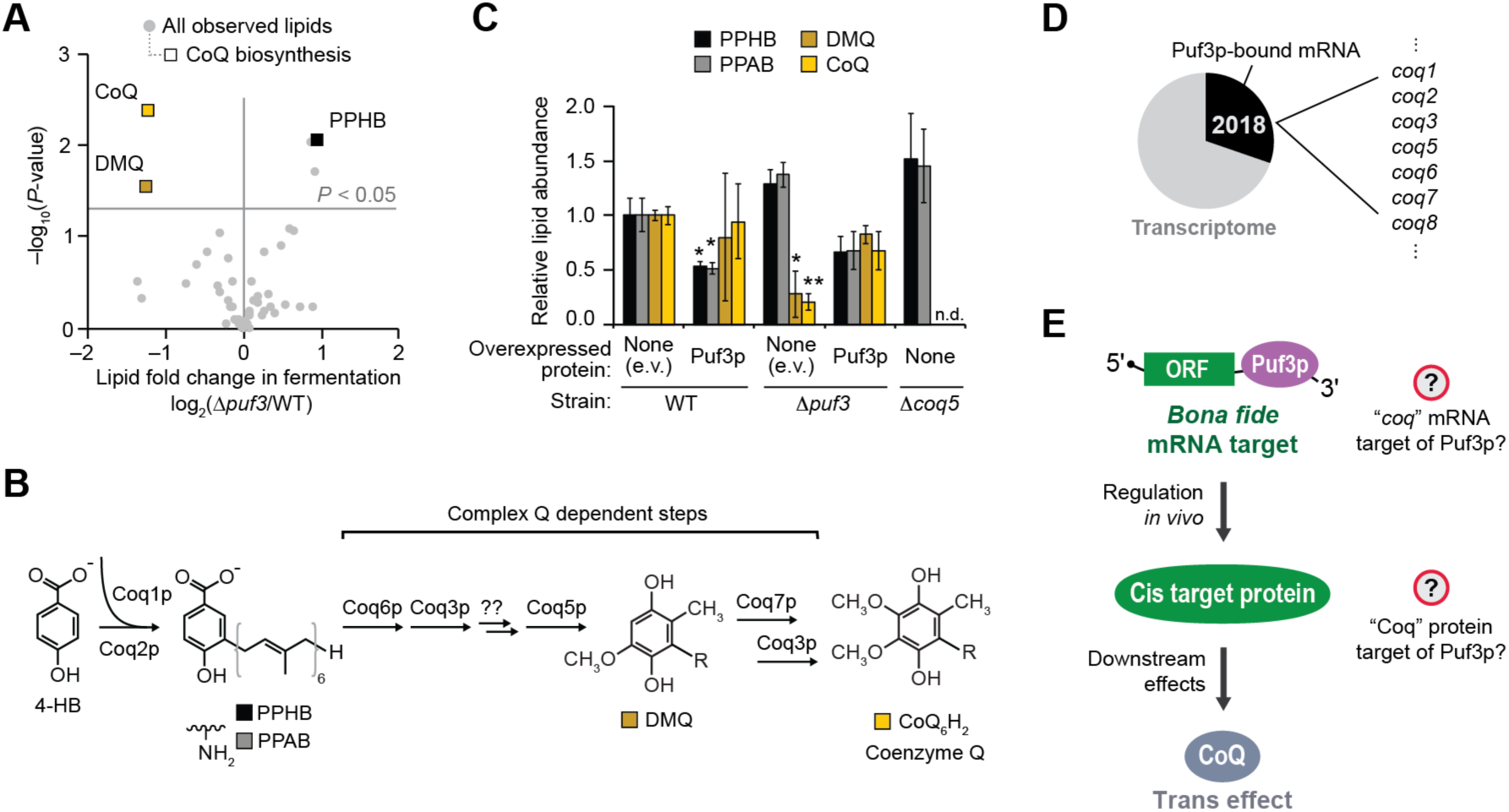
*puf3* Regulates Coenzyme Q Biosynthesis. (A) Relative lipid abundances in Δ*puf3* yeast compared to WT (mean, n = 3) (fermentation condition) versus statistical significance. Raw data from Y3K data set (Stefely et al., 2016a). (B) Scheme of CoQ biosynthesis. 4-HB, 4-hydroxybenzoate; PPHB, polyprenylhydroxybenzoate; PPAB, polyprenylaminobenzoate; DMQ, demethoxy-CoQ. (C) Relative lipid abundances in yeast transformed with low copy plasmids overexpressing Puf3p (or empty vector, e.v.) and cultured in fermentation media (mean ± SD, n = 3). Bonferroni corrected *p < 0.05; **p < 0.01. Two-sided Student’s *t*-test for all panels. (D) Pie chart illustrating the fraction of transcribed genes with mRNAs that are putative Puf3p targets, which was derived by aggregating all Puf3p-bound mRNAs reported via HITS-CLIP (Wilinski et al., 2017), RNA Tagging (Lapointe et al., 2015), PAR-CLIP (Freeberg et al., 2013), and RIP-seq (Kershaw et al., 2015). Puf3p-bound mRNAs include 7 mRNAs encoding CoQ biosynthesis enzymes. (E)	Scheme of how Puf3p could impact CoQ production.

To test these observations, we overexpressed Puf3p in wild type (WT) or Δ*puf3* yeast and examined CoQ pathway lipids. Overexpression of Puf3p in WT yeast suppressed production of PPHB and PPAB (the aminated analog of PPHB)—striking effects because they are the inverse of those observed in Δ*puf3* yeast (Figures 1C and S1C–S1E). Furthermore, plasmid expression of Puf3p recovered CoQ biosynthesis in fermenting Δ*puf3* yeast (Figure 1C). Thus, Puf3p regulates CoQ biosynthesis. We sought to understand how it does so.

We hypothesized that Puf3p directly controls an enzyme in the CoQ biosynthesis pathway by binding to and regulating the mRNA encoding it. To explore candidate enzymes, we examined existing data sets from recent studies that used genome-wide technologies to identify mRNAs bound by Puf3p (Freeberg et al., 2013; Kershaw et al., 2015; Lapointe et al., 2015; Wilinski et al., 2017). In aggregate, they reported an astounding 2,018 mRNA putative “targets” of Puf3p—nearly a third of the yeast transcriptome (**Table S1**). Putative Puf3p targets include seven enzymes directly involved in CoQ biosynthesis: *coq1*, *coq2*, *coq3*, *coq5*, *coq6*, *coq7*, and *coq8* (Figure 1D). However, given the extensive length of the Puf3p-bound mRNA list, we suspected that Puf3p regulates only a fraction of its 2,018 putative targets *in vivo*— both for this specific case of Puf3p-mediated regulation of CoQ (Figure 1E) and for Puf3p functions in general.

### Transomic Analyses Define Puf3p Targets

To identify high-confidence Puf3p targets, we integrated data from five experimental methods across four omic planes (Figure 2A). We first identified Puf3p target mRNAs using data from two independent approaches: HITS-CLIP (Licatalosi et al., 2008), which uses UV-crosslinking and immunopurification of RBP-RNA complexes, and RNA Tagging (Lapointe et al., 2015), which employs an RBP-poly(U) polymerase fusion protein to covalently tag mRNAs bound *in vivo* with 3’ uridines (“U-tags”) (Figure S2A). We focused on these datasets because they were generated in-house and rely on orthogonal strategies to identify mRNAs bound by a protein. HITS-CLIP (Wilinski et al., 2017) and RNA Tagging (Lapointe et al., 2015) co-identified 269 Puf3p-bound mRNAs, out of the 467 and 476 mRNAs identified via each technique on its own, respectively (Figure 2A) (hypergeometric test, p < 10^‒211^). Separating Puf3p-bound mRNAs into classes based on detection (I–IV, with class I representing the strongest detection) showed that co-identified mRNAs were the most robustly detected mRNAs in each method alone (Figures 2B, S2B, and S2C), and 42% of mRNAs identified via both methods were class I or II in each individual approach (Figure S2D). Moreover, the 3’ untranslated regions (UTRs) of mRNAs identified via both methods were strongly enriched for sequences that conform to high-affinity PBEs (UGUAHAUA), and a remarkable 81% also contained an upstream (–1 or –2) cytosine, which is required for particularly high-affinity interactions and regulation *in vivo* (Lapointe et al., 2015; Zhu et al., 2009) (Figures 2C and S2E). In contrast, RNAs unique to one method had more degenerate PBEs, with much less frequent upstream cytosines.

**Figure 2.**
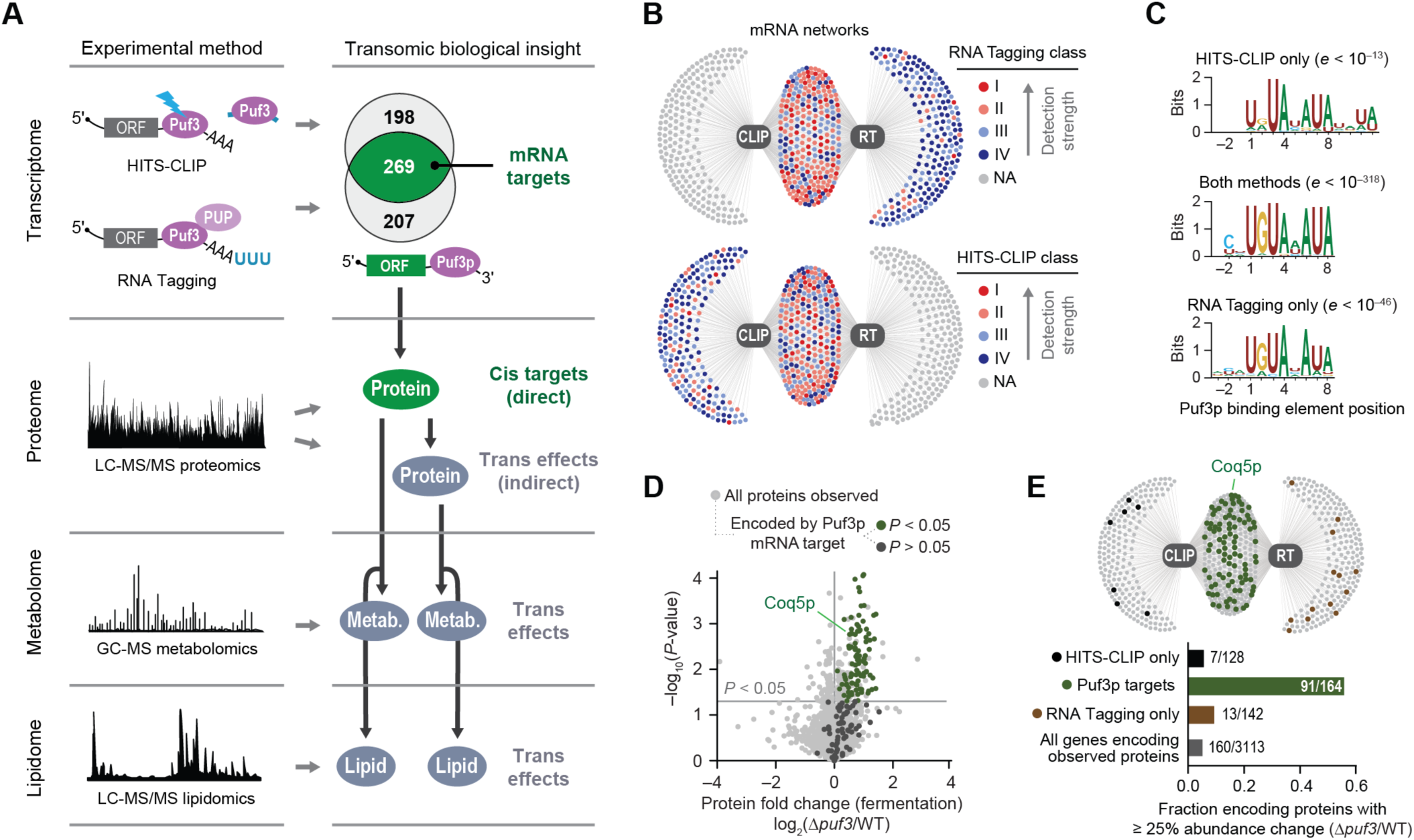
Transomics Maps High-Confidence Puf3p Targets. (A) Transomics approach overview. The Venn diagram indicates Puf3p-bound mRNAs co-identified by RNA Tagging and HITS-CLIP. (B) Network maps of Puf3p-bound mRNAs (dots) detected by RNA Tagging (RT) and/or HITS-CLIP (CLIP) (edges). (C) Enriched Puf3p-binding elements identified by MEME for the indicated groups of Puf3p-bound RNAs. (D)	Relative protein abundances in Δ*puf3* yeast compared to WT (mean, n = 3) versus statistical significance (fermentation condition), highlighting proteins encoded by Puf3p mRNA targets. *P*-value cutoff is for protein abundance changes. (E)	Bar graph and network map of Puf3p-bound mRNAs indicating mRNAs encoding proteins detected in the Y3K proteomics data set with ³ 25% protein abundance change (p < 0.05, two-sided Student’s *t*-test). This figure includes new, integrated analyses of publicly available raw data from the RT (Lapointe et al., 2015), HITS-CLIP (Wilinski et al., 2017), and Y3K multi-omic (Stefely et al., 2016a) data sets generated in our labs.

Integration of HITS-CLIP and RNA Tagging data thus yielded a high-confidence list of co-identified Puf3p-bound mRNAs, referred to here as “Puf3p target mRNAs” (Figure 2A and **Table S1**). Importantly, our analyses identified the CoQ-related mRNAs *coq2*, *coq5*, and *coq6* as high-confidence Puf3p targets, but not *coq1, coq3*, *coq7*, or *coq8*. In searching for the target responsible for the Δ*puf3* CoQ deficiency (Figure 1A), the observation of elevated PPHB and decreased CoQ suggested a defect in a complex Q dependent step (Figure 1B) and allowed us to narrow our focus to *coq5* and *coq6*. However, which of these Puf3p-mRNA interactions ultimately leads to regulation *in vivo* is unclear from the transcriptomic data alone.

To determine how Puf3p impacts the abundance of proteins encoded by its target mRNAs, we integrated these mRNA data with the proteomics feature of the Y3K data set (Stefely et al., 2016a). We identified protein abundance changes due to loss of Puf3p in two metabolic conditions. In fermentation culture conditions, 160 significant proteome changes were observed in Δ*puf3* yeast (p < 0.05 and fold change [FC] > 25%) (Figure S2F). In contrast, only 24 such changes in protein abundance were observed in respiration culture conditions. Thus, we primarily leveraged the fermentation proteomics data set to reveal Puf3p functions. The more drastic protein changes observed in fermentation mirror those of CoQ pathway lipids, which were likewise significantly (p < 0.05) altered only in fermentation (Figures 1A and S1B).

Of the 165 proteins encoded by Puf3p target mRNAs detected in fermenting yeast, 91 (55%) were significantly more abundant by at least 25% in Δ*puf3* yeast relative to WT yeast (p < 0.05) (Figure 2D). We collectively refer to these proteins as “Puf3p cis target proteins” (“cis targets”) (Figure 2A), and they include Coq5p but not Coq6p. The 91 cis targets account for a striking 57% (91/160) of significantly (p < 0.05) altered proteins in the Δ*puf3* proteome. In contrast, Puf3p-bound RNAs uniquely identified by a single RNA method, especially those detected weakly, were much less likely to be upregulated (Figures 2E and S2G). Thus, the combination of these distinct “omic” approaches identifies RNA-binding events likely to have regulatory effects in the cell. Collectively, these analyses provide high-confidence Puf3p targets across both the transcriptome and the proteome (Figure 2A and **Table S2**). They also demonstrate that Puf3p directly regulates Coq5p, suggesting a mechanistic link between loss of Puf3p and dysregulation of CoQ pathway lipids.

### Puf3p Binding to *coq5* mRNA Prevents Toxicity

To test the idea that Puf3p-mediated regulation of Coq5p abundance is required for proper CoQ production, we analyzed how Coq5p overexpression (i.e., overriding Puf3p regulation) impacted yeast growth and the CoQ biosynthetic pathway. Noticeably reduced CoQ levels can be sufficient for supporting respiratory yeast growth (Stefely et al., 2015), and alterations in CoQ biosynthesis can be difficult to track with growth assays. Thus, to facilitate mechanistic studies, we overexpressed Coq5p from a plasmid with a strong promoter to make associated phenotypes more readily observable. Overexpression of Coq5p in WT yeast slowed fermentative growth and essentially eliminated respiratory growth (Figures 3A and 3B). However, overexpression of Coq8p or Coq9p, which are also complex Q members (Floyd et al., 2016; He et al., 2014; Stefely et al., 2016b), did not inhibit yeast growth, supporting the hypothesis that regulation of Coq5p is particularly important for complex Q function (Figures 3A, 3B, and S3A). To examine whether Coq5p overexpression inhibits respiratory yeast growth by disrupting CoQ production, we examined CoQ pathway intermediates. Indeed, Coq5p overexpression in WT yeast recapitulated the CoQ-related phenotypes of Δ*puf3* yeast ― deficiency of DMQ and CoQ, and elevation of PPAB and PPHB (Figures 3C, S3B, and S3C). Similar to the observed growth effects, CoQ intermediates were most affected by overexpression of Coq5p compared to other proteins.

**Figure 3.**
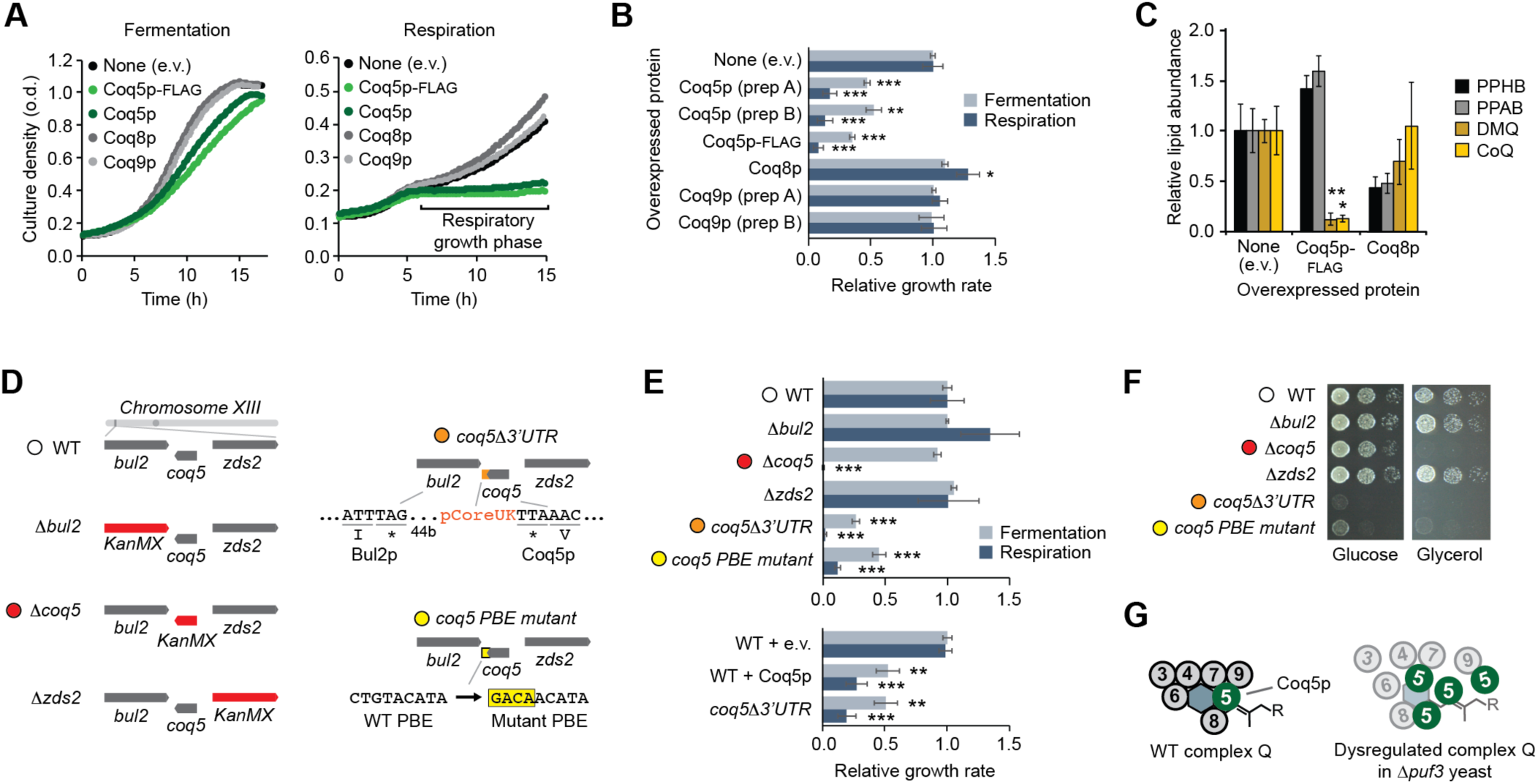
Puf3p Regulates the Potentially Poisonous CoQ Biosynthesis Enzyme Coq5p. (A) Growth curves for WT yeast transformed with plasmids overexpressing the proteins shown and cultured in either fermentation or respiration media. (B) Relative growth rates of WT yeast transformed with plasmids overexpressing the proteins shown and cultured in either fermentation or respiration media (mean ± SD, n = 3). **p < 0.01; ***p < 0.001. (C) Relative lipid abundances in WT yeast transformed with plasmids overexpressing the proteins shown and cultured in respiration media (mean ± SD, n = 3). Bonferroni corrected *p < 0.05; **p < 0.01. (D) Scheme of genetic alterations in the yeast strains used in (E) and (F). (E)	Yeast strain growth rates in either fermentation or respiration media (mean ± SD, n = 3). **p < 0.01; ***p < 0.001. (F)	Serial dilutions of yeast strains cultured on solid media containing either glucose or glycerol. (G)	Model for how Coq5p overexpression dysregulates complex Q. Two-sided Student’s *t*-test for all panels.

Coq5p overexpression could be deleterious due to excessive Coq5p methyltransferase activity or inappropriate Coq5p physical interactions. Overexpression of Coq5p with mutations to putative catalytic residues (D148A or R201A) inhibited yeast growth (Figure S3D), which suggests inhibition occurs through inappropriate physical interactions rather than excessive Coq5p enzyme activity. Furthermore, overexpression of Coq5p lacking its mitochondrial targeting sequence (MTS) (Coq5p^−MTS^) reduced its inhibitory effect in respiration (Figure S3E), suggesting that overexpressed Coq5p has inappropriate interactors in the mitochondrial matrix.

To further test the idea that Puf3p directly regulates Coq5p, we generated yeast strains with genomic mutations to the 3’ UTR of *coq5* (Figure 3D). Removal of the *coq5* 3’ UTR or the site-specific mutation of the Puf3p-binding element (PBE) in *coq5* at its endogenous genomic loci significantly reduced yeast growth in both fermentation and respiration (p < 0.05) (Figures 3E, 3F, and S3F). These growth phenotypes recapitulate those observed when Coq5p is overexpressed from a plasmid, but are notably different from those of Δ*coq5* yeast, which grow normally in fermentation (Figures 3E and 3F). As a control, disruption of two genes that flank *coq5* (*bul2* and *zds2*) had no discernible effect on yeast growth, therefore ensuring that the growth defects we observed are specific to loss of the PBE in *coq5*.

Our collective findings demonstrate that Puf3p modulates CoQ biosynthesis by directly regulating Coq5p, a potentially promiscuous protein with toxic effects when overexpressed (Figures 3G and S3G). By validating Coq5p as a Puf3p target, our findings strongly suggest that the additional 90 cis target proteins identified by our transomic analysis are also very likely to be *bona fide* Puf3p targets *in vivo.* They thus provide a foundation for identifying additional pathways that Puf3p coordinates with CoQ biosynthesis.

### Puf3p Coordinates CoQ Production with Mitochondrial Biogenesis Functions

Our transomic data set enabled us to map additional Puf3p functions across multiple omic planes (Figure 2A), similar to how we mapped Puf3p to its mRNA target *coq5*, its cis target protein Coq5p, and its downstream (trans) effect on CoQ lipids. We used the cis Puf3p protein targets to identify downstream “trans effects” in the proteome. By a protein “trans effect,” we refer to proteins whose abundance is dependent on Puf3p, but whose mRNA does not bind Puf3p (see Methods for details). The Δ*puf3* proteome changes include the 91 cis targets, which are enriched for mitochondrial organization and translation functions, and 49 trans effects, enriched for mitochondrial OxPhos and electron transport chain (ETC) functions (Figures 4A and S4A and **Tables S2** and **S3**).

We quantitatively compared properties of Puf3p cis targets and trans effects to those of either all proteins or mitochondrial proteins. The latter is an informative control set because Puf3p target mRNAs are highly enriched for mitochondrial proteins (Figure S4B). Both cis targets and trans effect proteins have relatively low abundances in fermentation (Stefely et al., 2016a), but not in respiration, reflecting Puf3p-mediated repression in fermentation (Figures 4B and S4C). In parallel, cis targets are upregulated across early time points in the diauxic shift (the metabolic shift from fermentation to respiration) (Stefely et al., 2016b) (Figure S4D), and yeast with respiration deficient mitochondria, which cannot complete the diauxic shift, exhibit downregulation of Puf3p cis targets as part of their “respiration deficiency response” (Stefely et al., 2016a) (Figure S4E). Cis targets are enriched for proteins that are co-translationally targeted to mitochondria (Williams et al., 2014) (Figures 4C and S4F). Furthermore, they are enriched for proteins that are toxic when overexpressed (Figure 4D), suggesting that regulation of toxic protein expression— such as that observed with Coq5p—is a general mechanism of Puf3p action. Together, our analyses support a model where Puf3p mediates repression of a set of mitochondrial proteins in fermentation, and this repression is released, or potentially reversed (i.e., cis targets are activated), early during the diauxic shift to help initiate the requisite biological changes, such as mitochondrial biogenesis (**Figure 4E**).

**Figure 4.**
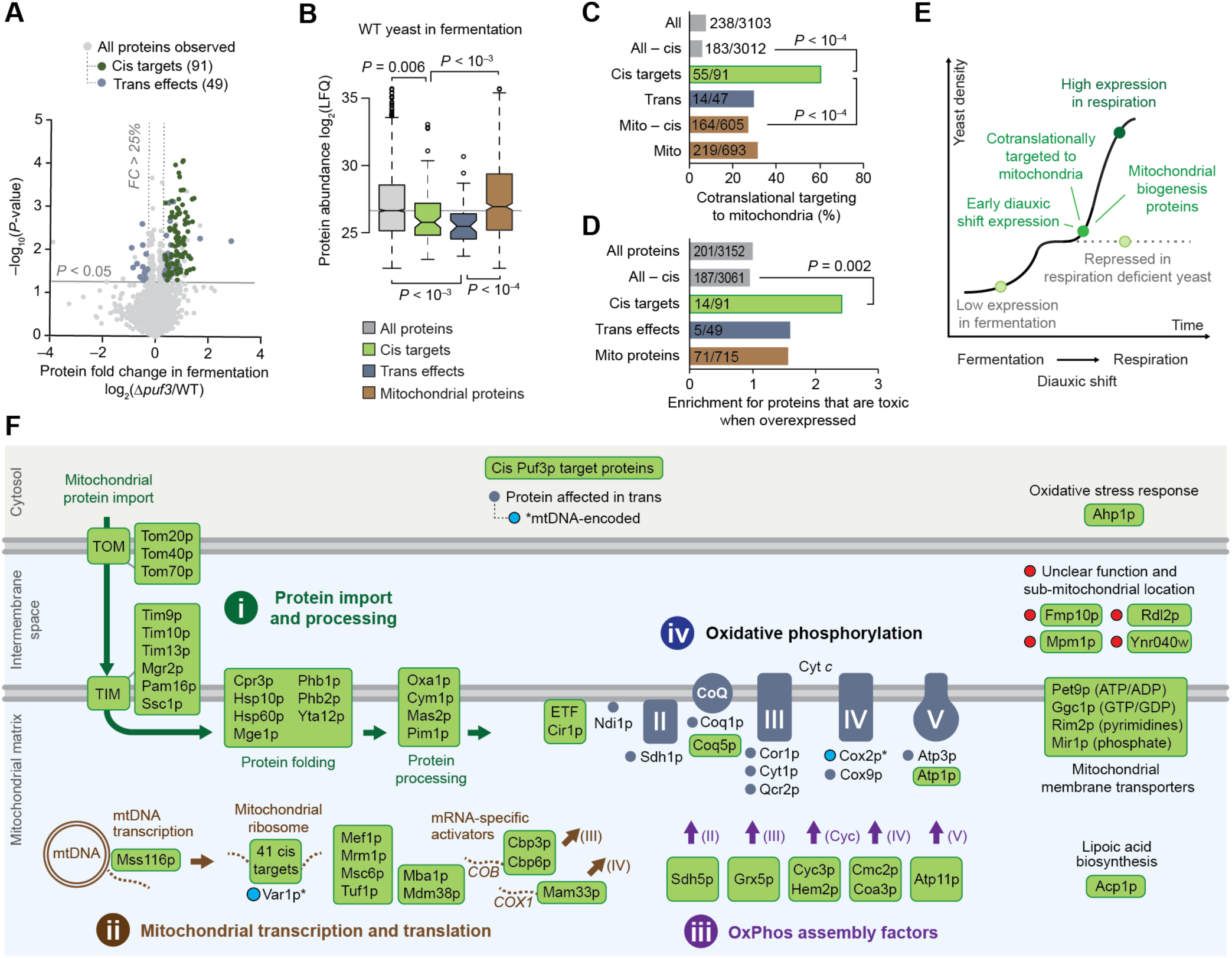
Puf3p Targets Prime Biogenesis of Mitochondria and OxPhos. (A) Relative protein abundances in Δ*puf3* yeast compared to WT (mean, n = 3) versus statistical significance, highlighting proteins with fold change (FC) > 25% and p < 0.05 (two-sided Student’s *t*-test) that were either identified as Puf3p targets by both RNA methods (cis targets, green) or neither RNA method (trans effects, blue). (B) Protein abundances (Stefely et al., 2016a) for each group of proteins shown. LFQ, label free quantitation value. Center lines indicate medians, limits indicate 25^th^ and 75^th^ percentiles, whiskers extend 1.5 times the interquartile range, outliers are represented by dots, and *P*-values were determined with a Student’s *t*-test (two-tailed, homostatic). Protein set sizes: all proteins n = 3152, mitochondrial proteins n = 715, cis targets n = 91, trans effect proteins n = 49. (C) Percent of each group of proteins shown that is cotranslationally targeted to mitochondria (Williams et al., 2014). *P*-values determined by a Fisher’s exact test. (D) Relative fraction of proteins that are toxic when overexpressed across the indicated protein groups. *P*-values determined by a Fisher’s exact test. (E)	Cartoon model of a yeast growth curve with key features and dynamics of Puf3p cis targets indicated. (F)	Cartoon map of all identified 91 Puf3p cis targets and select Puf3p trans effect proteins that are elevated in Δ*puf3* yeast. Puf3p cis targets include numerous proteins that support (i) mitochondrial protein import and processing, (ii) mitochondrial transcription and translation, and (iii) OxPhos assembly. Trans effects include (iv) an increase in OxPhos proteins.

To reveal the specific biochemical pathways through which Puf3p regulates mitochondrial function, we mapped individual Puf3p cis targets and trans effects (Figures 4F and S5A). Strikingly, 86 of the 87 Puf3p cis targets with known functions fit into pathways that support mitochondrial biogenesis, and in particular converge to generate the OxPhos machinery. The first pathway (i) includes proteins that catalyze the import, folding, and processing of nuclear DNA-encoded mitochondrial proteins, which include many OxPhos subunits. The second pathway (ii) includes proteins that support transcription and translation of mtDNA-encoded genes, which also encode OxPhos subunits. For example, cis Puf3p targets include over half of the mitochondrial ribosomal proteins and critical translational activators (e.g., Cbp3p, Cbp6p, Mam33p, Mba1p, and Mdm38p) (Figures 4F and S5A–S5C). Puf3p trans effects included increased abundance of two proteins encoded by mitochondrial DNA (Var1p and Cox2p, the only two such proteins observed). The third pathway (iii) encompasses assembly factors for each OxPhos subunit, which provide ancillary support for OxPhos complex biogenesis. Collectively, these three pathways of Puf3p cis targets are poised to prime OxPhos biogenesis. Consistently, Puf3p trans effects observed in Δ*puf3* yeast include increased abundance of OxPhos complex subunits (iv). Thus, Puf3p regulates production and assembly of both proteins and lipids required for OxPhos.

Two additional pathways of Puf3p cis targets include mitochondrial membrane transporters for nucleotides, which sustain mtDNA transcription, and proteins such as Acp1p that support the TCA cycle (Figure 4F). Downstream, citrate synthase (Cit1p), the rate limiting gateway enzyme to the TCA cycle and a putative client protein of the Puf3p cis targets Tom70p and Hsp60p (Martin et al., 1992; Yamamoto et al., 2009), and Ptc7p, a phosphatase that reactivates phosphorylated Cit1p (Guo et al., 2017), were also significantly increased trans effect proteins (Figure S6A). Accordingly, the abundance of citrate was significantly elevated in Δ*puf3* yeast (p < 0.05) (Figure S6B).

The tight functional association of cis Puf3p targets in mitochondrial biogenesis pathways (Figure 4F) suggests that the four uncharacterized proteins that are cis Puf3p targets ― Rdl2p, Ynr040w, Mpm1p, and Fmp10p ― might also function in mitochondrial biogenesis. Previous large-scale screens detected these proteins in mitochondria (Inadome et al., 2001; Reinders et al., 2006; Sickmann et al., 2003; Vogtle et al., 2012). Immunocytochemistry analyses of FLAG-tagged Rdl2p, Ynr040w, Mpm1p, and Fmp10p, validated these observations (Figures S7A and S7B), thereby confirming their classification as mitochondrial uncharacterized (x) proteins (MXPs). While the gene deletion strains for these MXPs are respiration competent, their overexpression in WT yeast inhibited respiratory growth (Figures S7C and S7D), suggesting that they may interact with proteins required for OxPhos and have the potential to be toxic like Coq5p. Given these data, all but one (Ahp1p) of the 91 cis targets localize to mitochondria, further solidifying the role of Puf3p in mitochondrial function. Together, this transomic analysis provides a high-7 resolution map of the molecular targets of Puf3p that will serve as a powerful resource for investigations of mitochondrial biogenesis and function (**Tables S1** and **S2**).

## DISCUSSION

### Defining Genuine Targets *via* Transomics

Our findings show that transomic strategies are powerful in the analysis of RBP function. In our case, despite considerable effort to identify the mRNAs to which Puf3p binds *in vivo*, it was unclear which of the many binding events were biologically productive – a central, common challenge in the analysis of RBPs (Konig et al., 2012; Licatalosi and Darnell, 2010; Riley and Steitz, 2013). To understand how Puf3p controlled CoQ biosynthesis, including whether there is a direct effect on an mRNA in the CoQ pathway, we generated a high-confidence Puf3p mRNA target list by combining two distinct approaches, and integrated that data with proteomic, metabolomic, and lipidomic studies. This strategy pinpointed Coq5p as the key Puf3p target from a large pool of candidates (> 2,000) that included six other CoQ biosynthesis enzymes (Figure S3G). Our analyses yielded a comprehensive roadmap of specific biochemical processes controlled by Puf3p *in vivo*. Together, our findings revealed a post-transcriptional mechanism to coordinate CoQ biosynthesis with the broader mitochondrial biogenesis program.

A key first step toward defining biochemical functions of Puf3p was the distillation of large lists of Puf3p-RNA interactions to a smaller set of those mRNAs it actually controls *in vivo* – its genuine targets *in vivo.* The HITS-CLIP and RNA Tagging data sets overlapped by ≈50%, which was greatest among the mRNAs detected most robustly in each separate approach. Weakly detected Puf3p-bound mRNAs were unlikely to overlap and likely arise through intrinsic biases of each method. Thus, our analyses illustrate that HITS-CLIP and RNA Tagging are complementary, and we demonstrate that integration of their data yields high-confidence RNA targets. Indeed, these analyses rapidly narrowed our focus to *coq5* or *coq6*, from the other complex Q candidates *coq3*, *coq8*, and *coq9*. Importantly, our findings also demonstrate that RBP-mRNA interaction data sets are stratified: interactions that lead to biological regulation are concentrated at the top of the rank order. In the absence of a transomic data set, a sharp focus on the top tier of candidates has great potential to reveal genuine targets.

A second key step was the integration of the Puf3p mRNA target set with Δ*puf3* proteome alterations. Our integration revealed 91 Puf3p-mRNA interactions that directly regulate the abundance of the encoded protein *in vivo*. The regulated Puf3p targets included Coq5p but not Coq6p, which yet again narrowed our focus and was critical to reveal how Puf3p controls CoQ biosynthesis. In principle, the remaining 74 Puf3p target mRNAs with unaffected protein abundances could be false positives. However, it is more likely that Puf3p regulates their localization (Gadir et al., 2011; Saint-Georges et al., 2008) independent of their translation, or that the yeast culture systems we used lacked an environmental stress encountered by yeast in nature. Notably, many of these mRNA targets trended toward increased protein abundance (Figure 2D), but did not cross our significance threshold (p < 0.05). They provide an initial foothold for analyzing alternative forms of Puf3p-mediated control (e.g., localization), or Puf3p function under varying environmental or cellular conditions.

### Post-Transcriptional Control of CoQ Biosynthesis and Toxic Proteins

Our results reveal a mechanism for post-transcriptional regulation of CoQ biosynthesis. By suppressing CoQ biosynthesis under fermentation conditions, Puf3p may help direct early CoQ precursors such as tyrosine and isoprene subunits to other biosynthetic pathways, such as protein synthesis and sterol biosynthesis, respectively.

Our biochemical work further demonstrated that Puf3p directly regulates the abundance of Coq5p, which when overexpressed dramatically reduced yeast growth and CoQ production. These findings suggest that the CoQ biosynthetic complex can be disrupted when particular subunits are too abundant, thereby highlighting the importance of stoichiometry for complex Q activity. Interestingly, however, overexpression of complex Q members Coq8p or Coq9p lacked the same deleterious effects as those observed with Coq5p, demonstrating that some complex Q subunits are particularly poisonous at inappropriately high protein abundance. Together, our findings suggest that Puf3p plays an important role in coordinating expression of complex Q components with the larger program of OxPhos biogenesis, through precise control of potentially toxic proteins. Analysis of the full Puf3p cis target set further suggested that regulation of poisonous proteins is a broader Puf3p function. This provides an evolutionary rationale for why particular proteins within larger pathways are targeted for regulation by Puf3p.

### Puf3p Coordinates Proteins and Lipids in Mitochondrial Biogenesis

By controlling the abundance of proteins that catalyze production of mitochondrial proteins, lipids, and metabolites, Puf3p regulation reaches across three omic planes. Transomic coordination may be critical for efficient production of functional mitochondria, for example by linking production of OxPhos protein complexes and the OxPhos lipid CoQ.

A similar mechanism was recently proposed for nuclear- and mitochondrial DNA-encoded proteins via synchronization of cytoplasmic and mitochondrial translation programs (Couvillion et al., 2016). That study suggested a critical role for cytosolic translational control of mitochondrial translational activators in coordinating OxPhos biogenesis and, in particular, identified the *COB* (complex III subunit) translational activator Cbp6p as a component of the mechanism. Importantly, our work now demonstrates that Puf3p directly regulates Cbp6p and its binding partner Cbp3p, both of which are cis Puf3p targets. Thus, we propose that Puf3p is a key part of the molecular mechanism that coordinates mitochondrial and cytosolic translation programs. Mammalian RBPs that impact mitochondrial function have been identified (Gao et al., 2014; Schatton et al., 2017), but whether any of these provide Puf3p-like transomic regulation of mitochondrial biogenesis is currently unknown.

### A Roadmap of Puf3p Functions

Our analyses provide substantial evidence that a primary function of Puf3p is to repress expression of mitochondrial biogenesis factors in fermenting yeast, as opposed to Puf3p having many broad biological functions (Kershaw et al., 2015). We identified 269 target mRNAs and 91 cis target proteins for Puf3p, which can be separated into three pathways that converge to generate the OxPhos machinery. Our strategy furthermore enabled us to map trans Puf3p effects across three omic planes, including protein components of OxPhos complexes, TCA cycle metabolites, and mitochondrial lipids, such as CoQ. Trans Puf3p effect proteins also include splicing factors and epigenetic regulators, and we speculate that Δ*puf3* yeast have “sensed” mitochondrial dysfunction through a mitochondria-to-nucleus retrograde signaling pathway. Consistently, the Y3K proteomics data set (Stefely et al., 2016a) shows that Δ*puf3* yeast have decreased levels of Ngg1p — a component of the SLIK complex, which regulates gene expression in response to mitochondrial dysfunction (Pray-Grant et al., 2002).

We observed few effects in respiring Δ*puf3* yeast, which at first glance may seem to contradict a recent study that reported Puf3p to activate translation of its mRNA targets as yeast enter the diauxic shift (Lee and Tu, 2015). However, our cultures were harvested more than 5 hours after the diauxic shift, by which time the activating function has disappeared (Lee and Tu, 2015). Indeed, our analyses of earlier time points also suggest that Puf3p functions early in the diauxic shift (Figure S4D). Yeast that lack *puf3* may only have mild phenotypes in respiration because transcriptional control compensates for the lack of Puf3p-mediated regulation, or because other RBPs (e.g., Puf4p or Puf5p) compensate for its absence. Regardless, Puf3p is a critical repressor of mitochondrial biogenesis in fermenting yeast ― a function that is conserved across more than 300 million years of evolution (Hogan et al., 2015; Wilinski et al., 2017).

The Puf3p mRNA targets that were not observed in our proteomics studies provide another valuable resource (**Tables S1** and **S2**). For example, they include previously uncharacterized genes (e.g., *aim11*, *aim18*, *ybr292c*, *ydr286c*, *ydr381c-a*, *ygr021w*, and *ygr161w-c*), which we can now potentially link to roles in mitochondrial biogenesis given their identity as Puf3p target mRNAs. Thus, the pathways mapped via transomics also provide a framework for retrospective analyses, which is particularly useful since imperfect overlap between omic planes is a common feature of transomic studies (e.g., some observed mRNAs are not detected at the protein level). Our high-confidence Puf3p cis target set provides a snapshot of specific mitochondrial biogenesis pathways regulated by Puf3p (Figure 4F) that can guide future analyses of the full set of Puf3p mRNA targets, which include additional OxPhos biogenesis factors.

Collectively, our work reveals post-transcriptional control of CoQ production linked into a larger regulatory network that controls OxPhos biogenesis. Our findings also provide a foundation for further exploring the molecular basis of mitochondrial biogenesis, including potential new roles for Puf3p-regulated MXPs in OxPhos biogenesis. Moreover, our approach presents a generalizable transomic strategy to identify biological roles for the many RNA-binding proteins that impact human health and disease.

## SUPPLEMENTAL FIGURES

**Figure S1, related to Figure 1.**
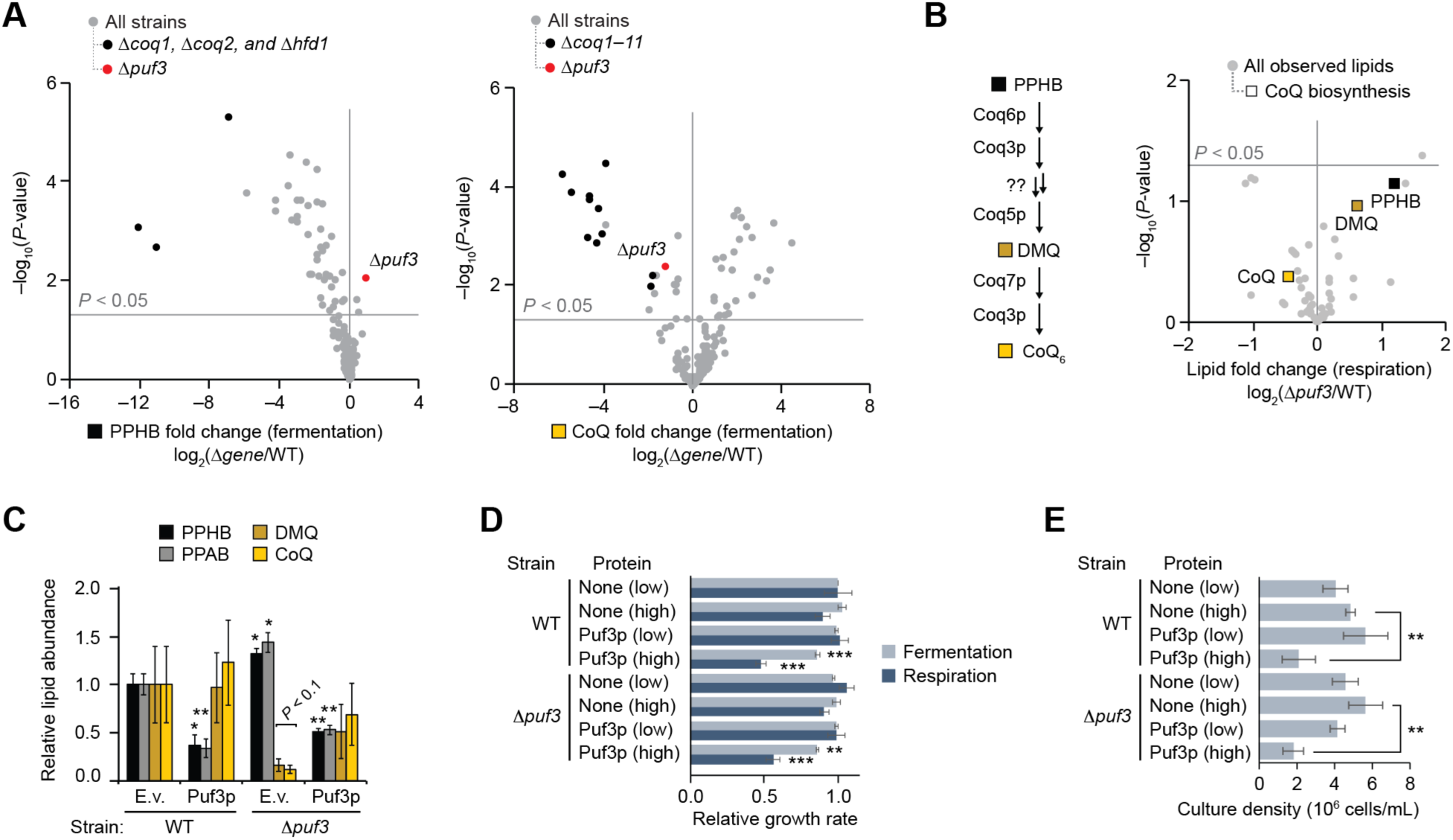
Puf3p Regulates CoQ Biosynthesis. (A) Relative abundances of PPHB and CoQ (mean, n = 3) versus statistical significance across all yeast strains in the Y3K data set (Stefely et al., 2016a) (fermentation culture condition). (B) Lipid abundances in Δ*puf3* yeast compared to WT (mean, n = 3) versus statistical significance (respiration condition). Raw data from the Y3K data set (Stefely et al., 2016a). (C) Relative lipid abundances in yeast transformed with high copy plasmids overexpressing Puf3p (or empty vector, e.v.) and cultured in fermentation media (mean ± SD, n = 3). Bonferroni corrected *p < 0.05; **p < 0.01. (D) Growth rates of WT or Δ*puf3* yeast transformed with plasmids overexpressing the proteins shown and cultured in either fermentation or respiration media (mean ± SD, n = 3). (E)	Culture densities of WT or Δ*puf3* yeast transformed with plasmids overexpressing the proteins shown and cultured in fermentation media (mean ± SD, n = 3) at the time point of harvest for the lipid analyses shown in Figure 1C and panel (C) of this figure. Two-sided Student’s *t*-test for all panels. *p < 0.05; **p < 0.01; ***p < 0.001

**Figure S2, related to Figure 2.**
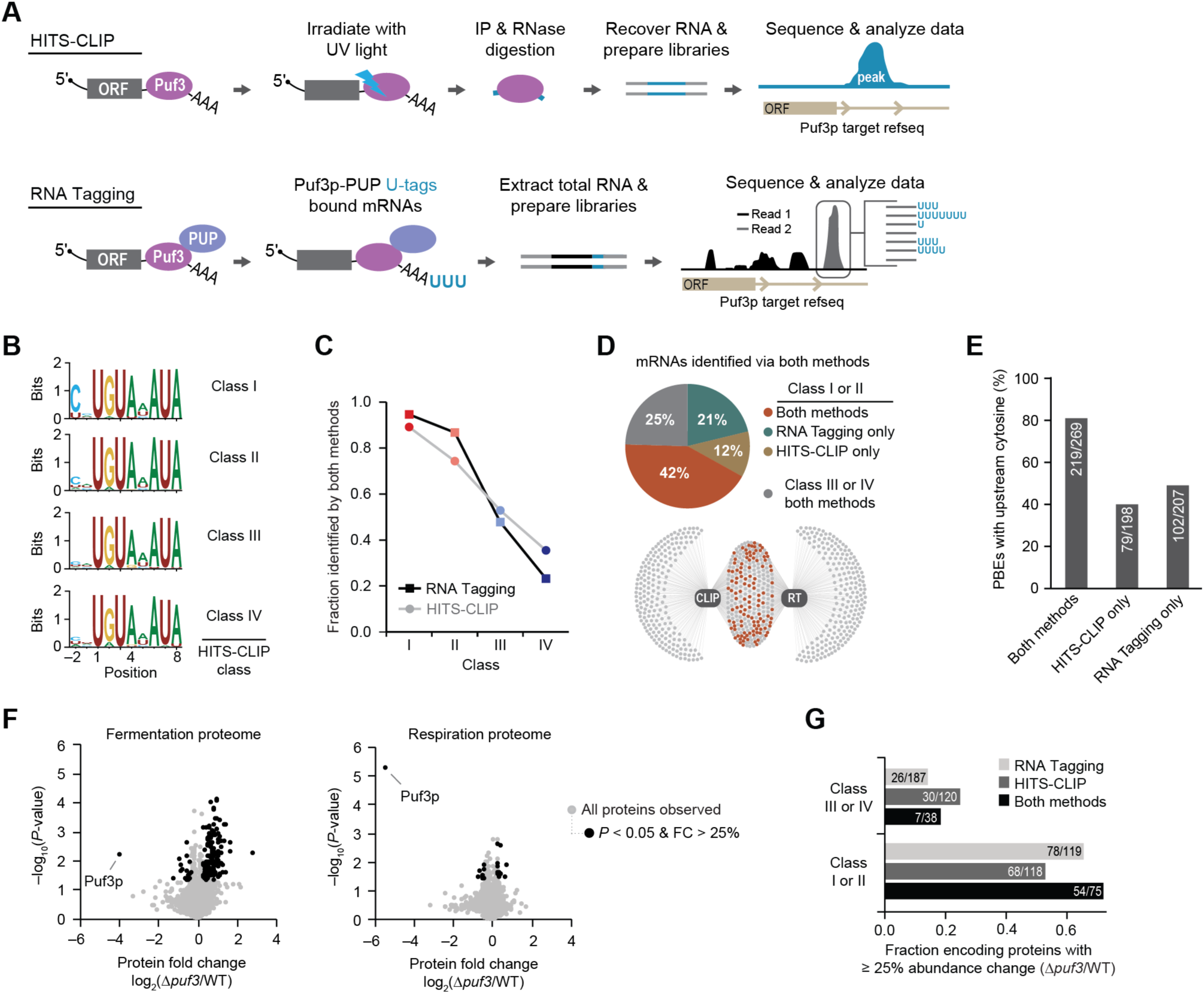
Integration of HITS-CLIP, RT, & Proteomics Defines Puf3p Targets. (A) Schematic of HITS-CLIP and RNA Tagging (RT) methods. (B) Position-weight matrices of PBEs identified under peaks for HITS-CLIP classes. (C) Overlap of RNAs identified as bound by Puf3p via both RNA Tagging and HITS-CLIP versus target class for the indicated method. (D) Class composition of Puf3p-bound mRNAs identified via both RNA Tagging (RT) and HITS-CLIP. The network map (bottom) shows Puf3p-bound mRNAs (dots) detected by RT and/or HITS-CLIP (CLIP) (edges). Puf3p-bound mRNAs present in class I or II of both methods are indicated in orange. (E)	Percent of PBEs with upstream cytosine (–1 or –2 position). (F)	Relative protein abundances in Δ*puf3* yeast compared to WT (mean, n = 3) versus statistical significance in fermentation and respiration conditions. Proteins with fold change (FC) > 25% and p < 0.05 (160 and 24 proteins, respectively) are highlighted (two-sided Student’s *t*-test). (G)	Fraction of genes with at least a 25% change in protein abundance (p < 0.05, two-sided Student’s *t*-test) for the indicated groups. Analyses were limited to genes with proteins detected in the Y3K proteomics study (Stefely et al., 2016a), which is the denominator of each ratio. This figure includes new, integrated analyses of publicly available raw data from the RT (Lapointe et al., 2015), HITS-CLIP (Wilinski et al., 2017), and Y3K multi-omic (Stefely et al., 2016a) data sets generated in our labs.

**Figure S3, related to Figure 3.**
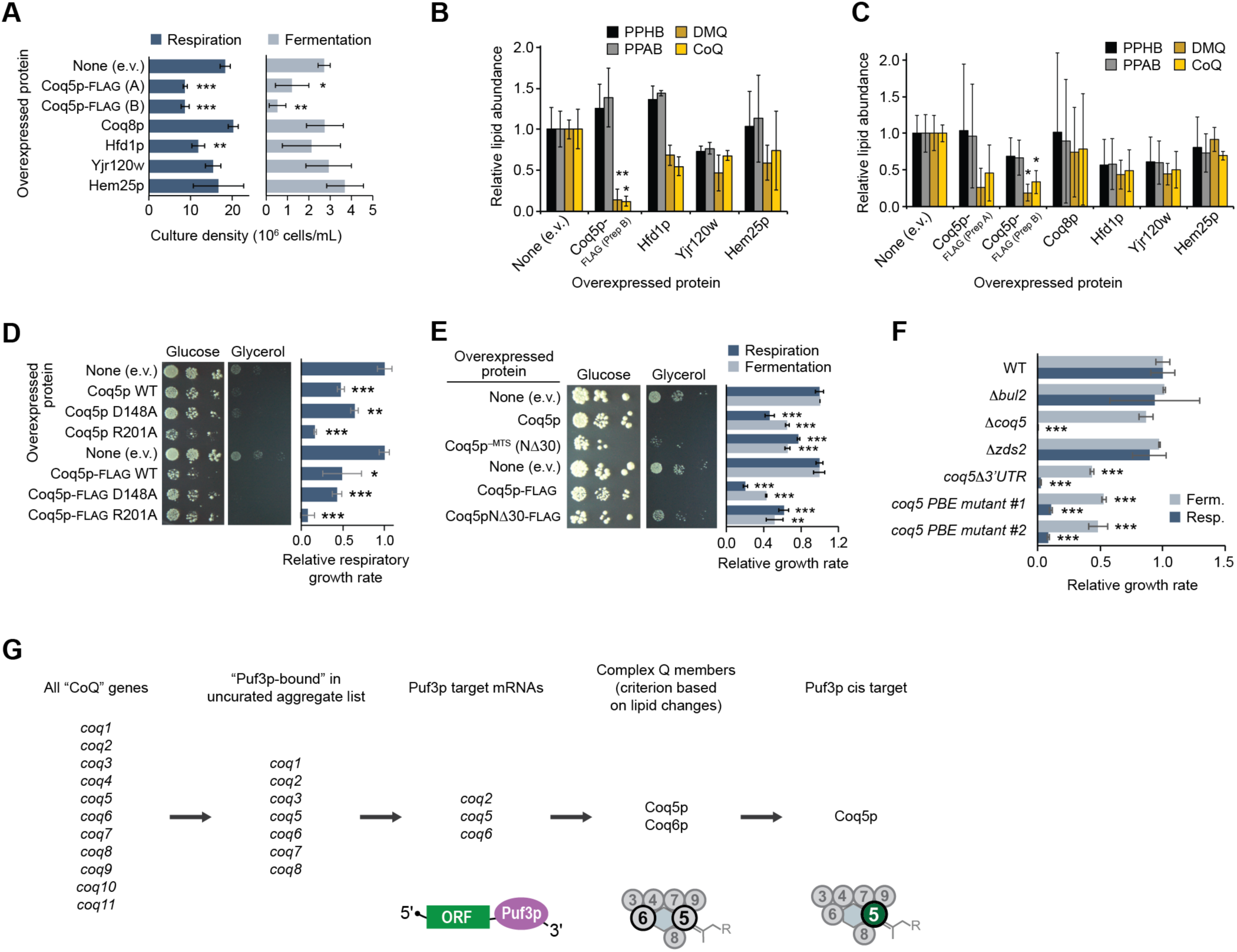
Puf3p Regulates the CoQ Biosynthesis Enzyme Coq5p. (A) Densities of yeast cultures at the time point of harvest for the lipid quantitation experiments depicted in Figure 3C and panels (B) and (C) of this figure. (B) and (C) Relative lipid abundances in WT yeast transformed with plasmids overexpressing the proteins shown and cultured in respiration media (B) or fermentation media (C) (mean ± SD, n = 3). Bonferroni corrected *p < 0.05; **p < 0.01 compared to empty vector (e.v.) control. (D) and (E) Left, serial dilutions of WT yeast transformed with plasmids overexpressing the proteins shown and cultured on solid media containing either glucose or glycerol. Right, growth rates of WT yeast transformed with plasmids overexpressing the proteins shown and cultured in liquid media (mean ± SD, n = 3). (F)	Growth rates of yeast strains cultured in either fermentation or respiration media (mean ± SD, n = 3). (G)	Scheme of how Coq5p was identified as a key Puf3p target responsible for the Δ*puf3* CoQ deficiency. For all panels, *P*-values were determined by a two-sided Student’s *t*-test. *p < 0.05; **p < 0.01; ***p < 0.001

**Figure S4, related to Figure 4.**
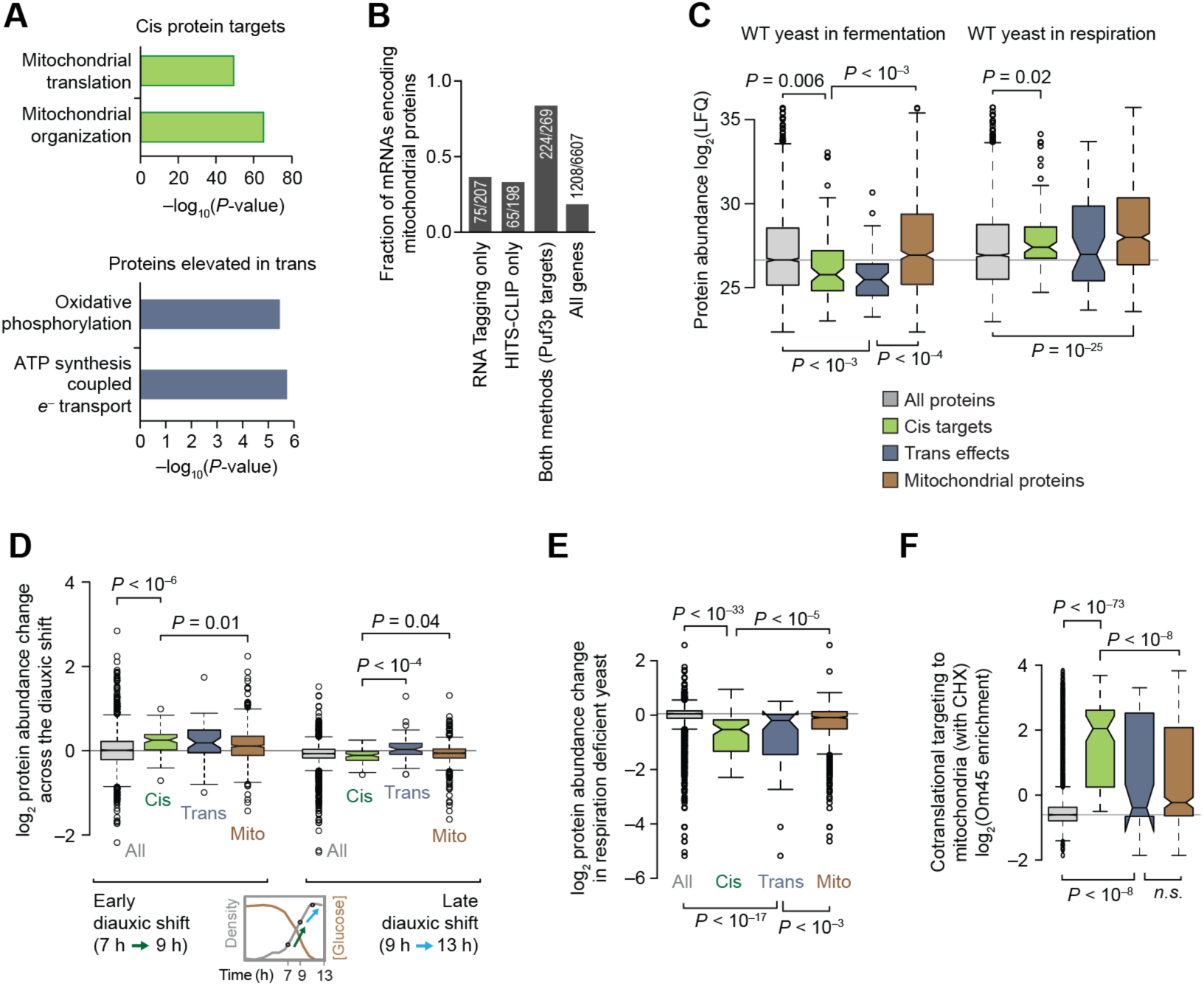
Puf3p Targets Share Distinctive Properties and Dynamics. (A) Gene Ontology (GO) terms enriched in cis Puf3p targets (top) or in elevated Puf3p trans effect proteins (bottom). (B) Fraction of genes annotated as encoding mitochondrial proteins for the indicated groups. (C)–(F) Box plots comparing the distributions of various gene and protein properties for all proteins observed in the Y3K Δ*puf3* proteomics data set (gray), cis Puf3p targets (green), trans Puf3p effects (blue), and mitochondrial proteins (brown). Center lines indicate medians, limits indicate 25^th^ and 75^th^ percentiles, whiskers extend 1.5 times the interquartile range, outliers are represented by dots, and *P*-values were determined with a Student’s *t*-test (two-tailed, homostatic). Protein set sizes: all proteins n = 3152, mitochondrial proteins n = 715, cis targets n = 91, trans effect proteins n = 49. (C) Protein abundances (Stefely et al., 2016a) for each group of proteins shown. LFQ, label free quantitation value. (D) Protein abundance changes (Stefely et al., 2016b) across the diauxic shift. (E)	Protein abundance changes (Stefely et al., 2016a) in respiration deficient yeast compared to respiration competent yeast. (F)	Measure of cotranslational targeting to mitochondria in yeast treated with cycloheximide (CHX) (Williams et al., 2014).

**Figure S5, related to Figure 4.**
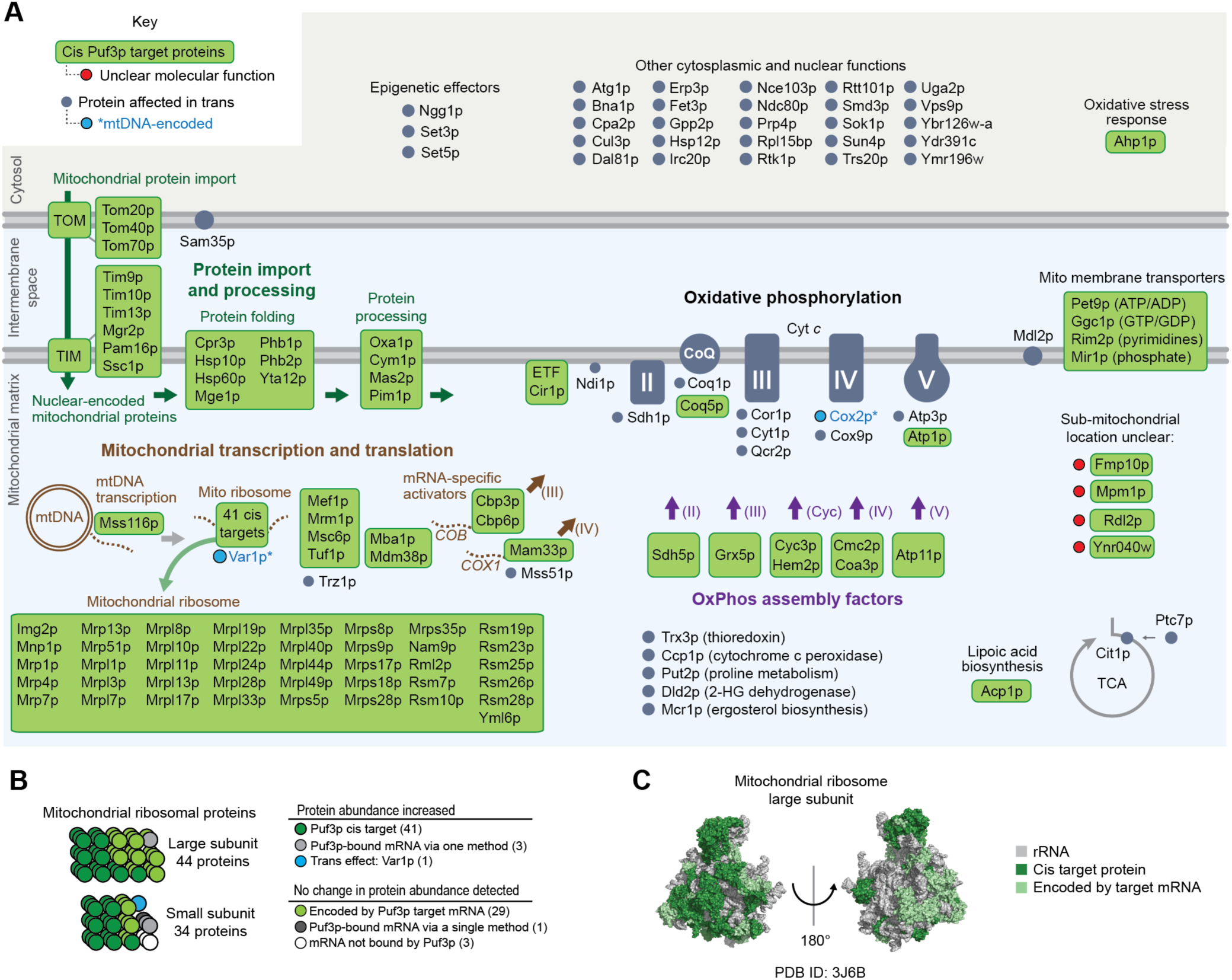
Puf3p Targets Pathways of Mitochondrial Biogenesis Proteins. (A) Cartoon indicating all identified cis Puf3p targets and trans effects. (B) Cartoon of mitochondrial ribosomal proteins with the effect by Puf3p indicated. (C) Surface representation of the large subunit of the yeast mitochondrial ribosome (PDB: 3J6B) (Amunts et al., 2014) indicating Puf3p targets.

**Figure S6, related to Figure 4.**
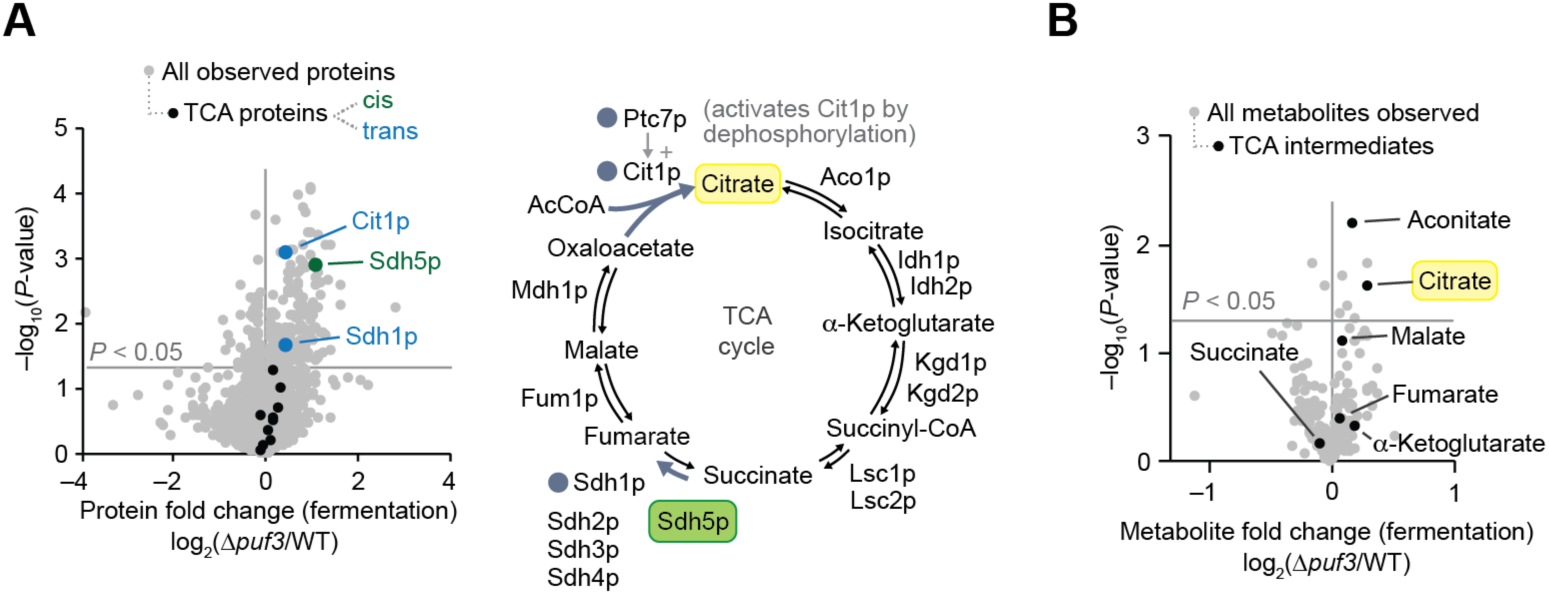
Puf3p Regulates TCA Cycle Proteins. (A) Relative protein abundances in Δ*puf3* yeast compared to WT highlighting TCA proteins (left), and scheme of the TCA cycle highlighting proteins significantly (p < 0.05) elevated in Δ*puf3* yeast (right). (B) Relative metabolite abundances in Δ*puf3* yeast compared to WT (mean, n = 3) versus statistical significance (fermentation condition), highlighting TCA cycle metabolites. Raw data from the Y3K data set (Stefely et al., 2016a).

**Figure S7, related to Figure 4.**
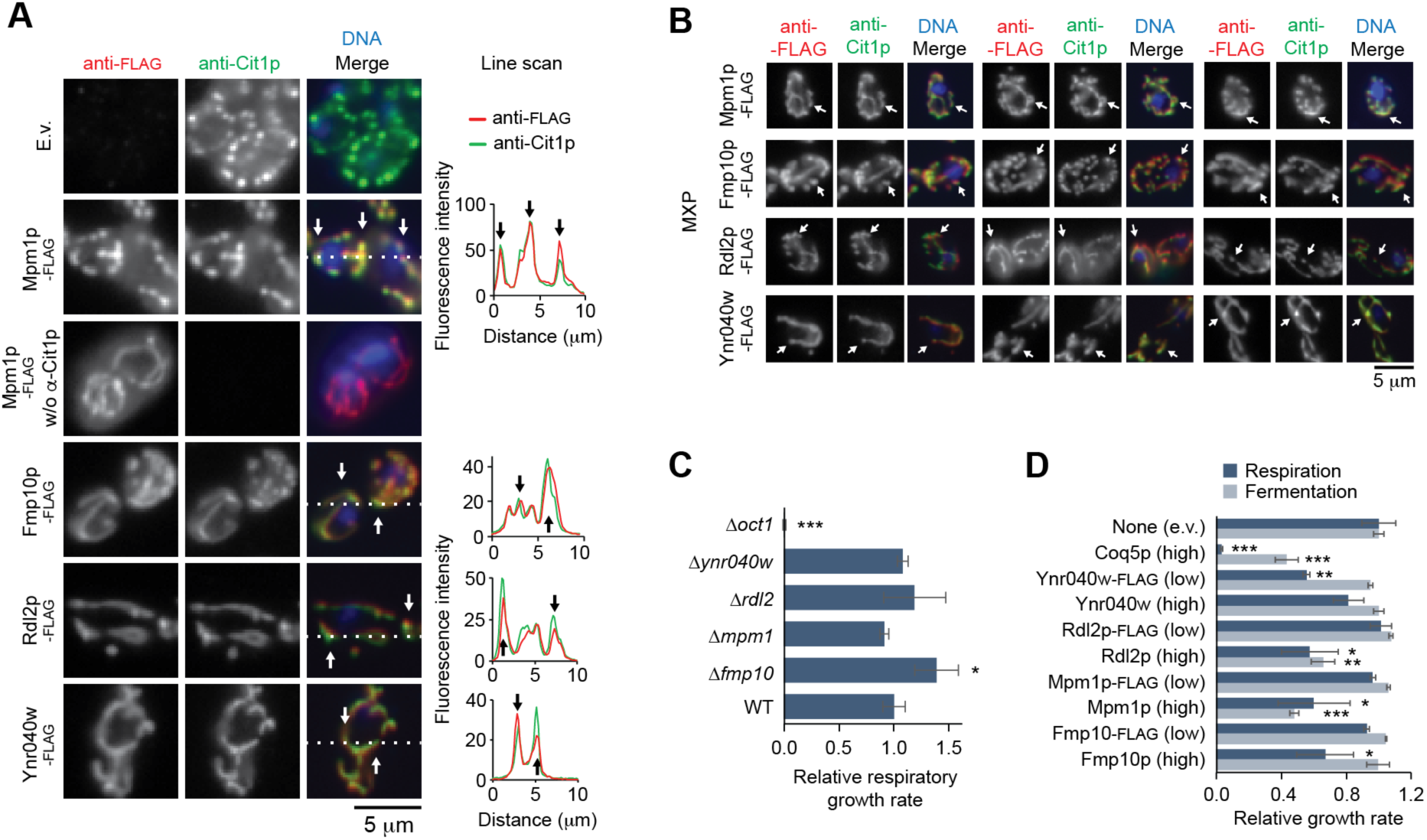
Uncharacterized Puf3p Targets Localize to Mitochondria. (A) Fluorescent immunocytochemistry analysis of yeast transformed with the indicated FLAG tagged proteins. An antibody against Cit1p was used to localize mitochondria. Arrows indicate landmarks on the line scan (dotted line on image). Scale bar, 5 µm. (B) Fluorescent immunocytochemistry analysis of WT yeast transformed with the indicated FLAG tagged mitochondrial uncharacterized proteins (MXPs). Arrows indicate spatial landmarks across each channel. Scale bar, 5 µm. (C) Relative growth rates of yeast strains cultured in respiration media (mean ± SD, n = 3). (D) Relative growth rates of WT yeast transformed with plasmids overexpressing the proteins shown and cultured in either fermentation or respiration media (mean ± SD, n = 3). High or low copy number plasmids are indicated. Two-sided Student’s *t*-test for all panels. *p < 0.05; **p < 0.01; ***p < 0.001

## SUPPLEMENTAL TABLE LEGENDS

**Table S1. Puf3p Target mRNA Analysis, Related to Figure 2**

(**A**) Key describing each column in sheets (tables) (B) – (D) of this data set: Gene systematic name, Gene standard name, RNA Tagging Class, Tagging U1 TRPM log_2_, Tagging PBE, CLIP peak height, CLIP *P*-value, CLIP Class, Number of peaks, Peak location, CLIP PBE, Method. (**B**) Table of Puf3p mRNA targets, including a summary of RNA Tagging and HITS-CLIP data for each target. (**C**) Table of all mRNAs identified as a Puf3p-bound mRNA in either approach, including a summary of RNA Tagging and HITS-CLIP data for each mRNA. (**D**) Table of all genes, highlighting mRNAs identified as a Puf3p-bound mRNA in either approach, including a summary of RNA Tagging and HITS-CLIP data for each mRNA. (**E**) Table of all genes, highlighting which method identified them as putative Puf3p targets (HITS-CLIP, RNA Tagging, PAR-CLIP, or RIP-seq). It includes a list of the aggregate 2,018 mRNAs identified as putative Puf3p targets.

**Table S2. Transomic Puf3p Target Analysis, Related to Figure 2**

(**A**) Key describing each column in sheets (tables) (B) – (D) of this data set: Gene systematic name, Gene standard name, Pathway, Protein effect (cis, trans, or unclear), RNA Tagging Class, Tagging U1 TRPM log_2_, CLIP peak height, CLIP *P*-value, CLIP Class, Number of peaks, Peak location, Method, Proteomics log_2_(KO/WT) in fermentation, Proteomics *P*-value in fermentation, and 25% sig protein change. (**B**) Table of Puf3p cis targets, including a summary of RNA Tagging, HITS-CLIP, and proteomics data for each target. (**C**) Table of Puf3p trans effect proteins, including a summary of the proteomics data for each protein. (**D**) Table of all genes and all proteins in the transomic analysis, including a summary of RNA Tagging, HITS-CLIP, and proteomics data for each molecule.

**Table S3. Puf3p Cis Target GO Term Analysis, Related to Figure S4**

(**A**) Table of Gene Ontology (GO) terms enriched in the Puf3p cis target set. Includes GO terms, *P*-values, –log_10_(*P*-values), genes in the Puf3p cis target set that overlap with those in the GO term set, and GO term ID number. (**B**) Table of Gene Ontology (GO) terms enriched in the set of trans effect proteins with increased protein abundance in yeast that lack Puf3p. Includes GO terms, *P*-values, –log_10_(*P*-values), genes in the Puf3p trans effect set that overlap with those in the GO term set, and GO term ID number.

## ACKNOWLEDGEMENTS

We thank members of the Pagliarini, Wickens, and Coon laboratories for helpful discussions. This work was supported by a UW2020 award and National Institutes of Health (NIH) grants R01GM112057 and R01GM115591 (to D.J.P.); NIH P41GM108538 (to J.J.C. and D.J.P.); NIH R01GM50942 (to M.P.W.); NIH R35GM118110 (to J.J.C.); Department of Energy Great Lakes Bioenergy Research Center (Office of Science BER DE-FC02-07ER64494 to N.W.K.); NIH fellowship T32GM008349 (to G.M.W.); Wharton and Biochemistry Scholar fellowships (to C.P.L.); and NIH Ruth L. Kirschstein NRSA F30AG043282 (to J.A.S.).

## AUTHOR CONTRIBUTIONS

C.P.L., J.A.S., M.P.W., and D.J.P. conceived of the project and its design and wrote the manuscript. J.A.S. and A.J. prepared samples and performed biochemical experiments. P.D.H. and G.W. acquired MS data. C.P.L., J.A.S., A.J., P.D.H., G.M.W., N.W.K., J.J.C., M.P.W., and D.J.P. analyzed data.

## METHODS

### CONTACT FOR REAGENT AND RESOURCE SHARING

Further information and requests for resources and reagents should be directed to and will be fulfilled by the Lead Contact David Pagliarini.

### EXPERIMENTAL MODEL AND SUBJECT DETAILS

#### Yeast strains

The parental (wild type, WT) *Saccharomyces cerevisiae* strain for this study was the haploid MATalpha BY4742. Single gene deletion (D*gene*) derivatives of BY4742 were obtained through the gene deletion consortium (via Thermo, Cat#YSC1054). Gene deletions were confirmed by proteomics (significant [p < 0.05] and selective decrease in the encoded protein) or PCR assay.

*Coq5 PBE mutant strain*. The PBE in *coq5* was identified by analysis of the HITS-CLIP data (Wilinski et al., 2017), in which *coq5* had a single peak centered over the sequence: CTGTACATA. Using the haploid MATalpha BY4742 as the parental strain, a yeast strain with a mutant Puf3p-binding element (PBE) in the 3’ untranslated region (UTR) of the *coq5* gene was generated using the *Delitto Perfetto* method (Storici and Resnick, 2006). Briefly, the 282 nucleotides immediately downstream of the *coq5* coding sequence were replaced with the KanMx4-KlUra3 cassette. Thus, the “*coq5*D3’UTR” strain was created. To enable generation of the endogenous PBE mutant, the *coq5* ORF + 3’ UTR was cloned in p426gpd. The PBE sequence was mutagenized by PIPE cloning. The sequence-confirmed plasmid, containing mutagenized PBE, served as the PCR template in a reaction that generated the cassette that seamlessly replaced KanMx4-KlUra3 with the *coq5* 3’ UTR sequence harboring the CTGT --> GACA PBE mutation. Two separate yeast strains with identical *coq5* PBE mutants were independently generated in this manner. One of the two strains maintained a *coq5* 3’ UTR identical to that of the WT strain outside of the mutated PBE. In the second strain, the PBE change was scarless, with the exception of the deletion of a single 3’ UTR thymidine nucleotide in a stretch of 10 thymidine nucleotides beginning 184 nucleotides upstream of the desired PBE mutation. As shown in the figures of this report, both of these independently generated strains shared the same biological phenotypes.

### METHOD DETAILS

#### Yeast cultures

*General culture procedures and media components*. Yeast were stored at −80 °C as glycerol stocks and initially cultured on selective solid media plates at 30 °C. Biological replicates were defined as separate yeast colonies after transformation and plating onto solid selective media. Experiments were conducted in biological triplicate (“n = 3”) to afford statistical power. Cell density of liquid media cultures was determined by optical density at 600 nm (OD_600_) as described (Hebert et al., 2013). Media components included yeast extract (‘Y’) (Research Products International, RPI), peptone (‘P’) (RPI), agar (Fisher), dextrose (‘D’) (RPI), glycerol (‘G’) (RPI), uracil drop out (Ura^−^) mix (US Biological), histidine drop out (His^−^) mix (US Biological), and G418 (RPI). YP and YPG solutions were sterilized by automated autoclave. G418 and dextrose were sterilized by filtration (0.22 µm pore size, VWR) and added separately to sterile YP or YPG. YPD+G418 plates contained yeast extract (10 g/L), peptone (20 g/L), agar (15 g/L), dextrose (20 g/L), and G418 (200 mg/L). YPD media (rich media fermentation cultures) contained yeast extract (10 g/L), peptone (20 g/L), and dextrose (20 g/L). YPGD media (rich media respiration cultures) contained yeast extract (10 g/L), peptone (20 g/L), glycerol (30 g/L) and dextrose (1 g/L). Synthetic Ura^−^ media were sterilized by filtration (0.22 µm pore size). Ura^−^,D media contained Ura^−^ mix (1.92 g/L), yeast nitrogen base (YNB) (6.7 g/L, with ammonium sulfate and without amino acids), and dextrose (20 g/L). Ura^−^,GD media contained Ura^−^ mix (1.92 g/L), YNB (6.7 g/L), glycerol (30 g/L), and dextrose (1 g/L). Ura^−^,D+4HB media contained Ura^−^ mix (1.92 g/L), YNB (6.7 g/L), dextrose (20 g/L), and 4-HB (100 µM). Ura^−^,GD+4HB media contained Ura^−^ mix (1.92 g/L), YNB (6.7 g/L), glycerol (30 g/L), dextrose (1 g/L), and 4-HB (100 µM).

*Puf3 rescue cultures.* WT or Δ*puf3* yeast were transformed with plasmids encoding Puf3p [p423(2µ)-PUF3 or p413(CEN)-PUF3] and cultured on His^−^,D plates. Starter cultures (3 mL His^−^,D+4HB) were inoculated with an individual colony of yeast and incubated (30 °C, 230 rpm, 10–15 h). For CoQ quantitation, His^−^,D+4HB (fermentation) or His^−^,GD+4HB (respiration) media (100 mL media at ambient temperature in a sterile 250 mL Erlenmeyer flask) was inoculated with 2.5×10^6^ yeast cells and incubated (30 °C, 230 rpm). Samples of the His^−^,D+4HB cultures were harvested 13 h after inoculation, a time point that corresponds to early fermentation (logarithmic) growth. Samples of His^−^,GD+4HB cultures were harvested 25 h after inoculation, a time point that corresponds to early respiration growth. For each growth condition, 1×10^8^ yeast cells were pelleted by centrifugation (3,000 *g*, 3 min, 4 °C), the supernatant was removed, and the cell pellet was flash frozen in N_2(l)_ and stored at -80 °C prior to lipid extractions. For relative growth rate measurements, analogous cultures (initial density of 5×10^6^ cells/mL) were incubated in a sterile 96 well plate with an optical, breathable cover seal (shaking at 1096 rpm). Optical density readings were obtained every 10 min.

*Protein overexpression cultures.* WT yeast were transformed with plasmids encoding Coq5p, Coq5p-FLAG, Coq8p, Coq9p, Hfd1p, Yjr120w, or Hem25p (p426[2µ]-GPD plasmids) and cultured on Ura^−^,D plates. Starter cultures (3 mL Ura^−^,D+4HB) were inoculated with an individual colony of yeast and incubated (30 °C, 230 rpm, 10–15 h). For CoQ quantitation, Ura^−^,D+4HB (fermentation) or Ura^−^,GD+4HB (respiration) media (100 mL media at ambient temperature in a sterile 250 mL Erlenmeyer flask) was inoculated with 2.5×10^6^ yeast cells and incubated (30 °C, 230 rpm). Samples of the Ura^−^,D+4HB cultures were harvested 13 h after inoculation. Samples of Ura^−^,GD+4HB cultures were harvested 25 h after inoculation. For each growth condition, 1×10^8^ yeast cells were pelleted by centrifugation (3,000 *g*, 3 min, 4 °C), the supernatant was removed, and the cell pellet was flash frozen in N_2(l)_ and stored at -80 °C prior to lipid extractions. For relative growth rate measurements, analogous cultures (initial density of 5×10^6^ cells/mL) were incubated in a sterile 96 well plate with an optical, breathable cover seal (shaking at 1096 rpm). Optical density readings were obtained every 10 min. Growth rates were determined by fitting a linear equation to the linear growth phase and determining the slope of the line.

*Cultures for microscopy*. BY4742 *S. cerevisiae* overexpressing C-terminally FLAG-tagged genes from p416gpd_FLAG plasmids were cultured in Ura^−^,D media (30 °C, 3 mL starter cultures, ~14 h). From these starter cultures, 1.25×10^6^ cells were used to inoculate Ura^−^,GD media (50 mL) (respiration culture condition). After incubating 25 hours (30 °C, 230 r.p.m), 1×10^8^ cells were removed from the culture by pipetting and immediately fixed with formaldehyde for microscopy as described below.

#### Yeast Transformations

Yeast were transformed with plasmids using a standard lithium acetate protocol (Gietz et al., 1992). Briefly, BY4742 yeast were cultured in YEPD (50 mL) to a density of 2×10^7^ cells/mL. Cells were pelleted and washed twice with water. For each transformation, added PEG 3350 (50% w/v, 240 µL), lithium acetate (1 M, 36 µL), boiled salmon sperm DNA (5 mg/mL, 50 µL), water (30 µL), and plasmid (4 µL) to a pellet containing 1×10^8^ cells, mixed by vortexing, and incubated (42 °C, 45 min). The transformed cells were pelleted, resuspended in water (100 µL), and plated on selective media.

#### DNA Constructs

Yeast gene constructs were generated by amplifying the *S. cerevisiae* genes *fmp10*, *mpm1*, *rdl2*, and *ynr040w* from strain BY4742 genomic DNA with primers containing HindIII recognition sequence (forward) and SalI recognition sequence (reverse). Similarly, *hem25* and *yjr120w* were amplified with BamHI (forward) and EcoRI (reverse) primers. *Coq5* (from strain W303) was amplified with SpeI (forward) and SalI or XhoI (reverse) primers. PCR reactions contained 1× Accuprime PCR mix, 1 µM forward primer, 1 µM reverse primer, ~250 ng template, and 1× Accuprime Pfx (Invitrogen cat#12344024). After an initial 2 min denaturation at 95 °C, reactions were exposed to 5 cycles of 95 °C for 15 seconds, 55 °C for 30 seconds, and 68 °C for 2 minutes followed by 30 cycles of 95 °C for 15 seconds, 60 °C for 30 seconds, and 68 °C for 2 minutes. Amplicons were purified using a PCR purification kit (Thermo cat#K0702) and digested with the appropriate restriction enzymes and again subjected to PCR purification. Amplified genes were cloned into restriction enzyme digested yeast expression vectors (p426gpd and/or p416gpd_FLAG). The plasmid p416gpd_FLAG was generated by digesting p416gpd with XhoI and MluI and inserting a double stranded oligonucleotide containing the Flag tag nucleotide sequence and processing XhoI and MluI ends. For *puf3*, p423(2µ)-PUF3 and p413(CEN)-PUF3 expression plasmids were constructed in modified p423(2µ) or p413(CEN) backgrounds in which the plasmid promoter and terminators had been removed. The *puf3* gene, including 1,000 upstream nts and 457 downstream nts, was amplified from BY4742 genomic DNA using a standard Phusion DNA polymerase PCR. Amplified products were cleaned via a PCR purification kit (ThermoFisher), digested with the SalI and KpnI restriction enzymes under standard conditions, and cloned into the appropriate plasmid. All recombinants were confirmed by DNA sequencing. Cloning was previously reported for p426-GPD-*coq8* (Stefely et al., 2015), p426-GPD-*coq9* (Lohman et al., 2014), p426-GPD-*hfd1* (Stefely et al., 2016a).

#### HITS-CLIP Class Definition

Puf3p-bound RNAs identified via HITS-CLIP (Wilinski et al., 2017) were sorted by the number of RNAs detected in their CLIP peak (“peak height”) from most to least. Classes were then defined as follows: the top 10% were designated class I; 11–40% class II; 41–70% class III; and 71–100% class IV. Classes were defined to be of comparable size to the analogous RNA Tagging class to facilitate cross-method comparisons (Lapointe et al.; Lapointe et al., 2015). As a control, the signal detected for these mRNAs [log_2_(TRPM) for RNA Tagging and log_2_(peak height) for HITS-CLIP] was correlated across methods (Spearman’s *ρ* = 0.42, p < 10^‒12^), but not with mRNA abundance (p > 0.01).

#### MEME Analyses

For all analyses, the 3ʹ untranslated regions (UTRs) was defined as the longest observed isoform for a gene (Xu et al., 2009) or 200 nts downstream of the stop codon if not previously defined. MEME was run on a local server with the command: meme [input.txt] -oc [outputdirectory] -dna -mod zoops -evt 0.01 -nmotifs 10 -minw 8 -maxw 12 -maxsize 100000000000.

#### Definition of Puf3p mRNA Targets

Puf3p HITS-CLIP data were obtained from (Wilinski et al., 2017). Only peaks that were assigned to annotated genes were considered: 467 in total, with 15 genes identified by two peaks. RNA Tagging data were obtained from (Lapointe et al.; Lapointe et al., 2015). All 476 reported RNA Tagging targets were considered. RNA Tagging classes were obtained from (Lapointe et al.) because they were generated using an improved strategy than in the initial report (Lapointe et al., 2015). Genes with mRNAs identified by both approaches, visualized by Venn diagrams, were designated as “Puf3p target mRNAs”. Protein-RNA network maps were constructed using Cytoscape (v. 3.2.1) and the ‘Organic’ ‘yFiles Layouts’ option.

#### Definition of Puf3p Cis Target Proteins

We defined “Puf3p cis target proteins” (interchangeably referred to as “Puf3p cis targets”) as: proteins encoded by Puf3p mRNA targets with at least a 25% significant (p < 0.05) alteration in protein abundance in yeast that lack *puf3* relative to WT yeast grown in fermentation culture conditions. The proteomic data included 165 proteins encoded by Puf3p mRNA targets (out of 269 total), and 91 proteins were designated as Puf3p cis target proteins (“Puf3p cis targets”) out of the 160 proteins with at least a 25% significant alteration in protein abundance. To ensure rigorous definition, we excluded 20 proteins with significantly altered protein abundances because they were identified as Puf3p-bound mRNAs only via a single method, thus confounding their assignment.

#### Definition of Puf3p Trans Targets

For the proteome, Puf3p trans effects were defined as proteins that were *not* encoded by Puf3p mRNA targets with at least a 25% significant (p < 0.05) alteration in protein abundance in yeast that lack *puf3* relative to WT yeast grown in fermentation culture conditions. For the metabolome and lipidome, Puf3p trans effects were defined as metabolites or lipids, respectively, with at least a 25% significant (p < 0.05) alteration in abundance in yeast that lack *puf3* relative to WT yeast grown in fermentation culture conditions.

#### Gene Ontology Analyses

Analyses were conducted using YeastMine, from the Saccharomyces Genome database (http://yeastmine.yeastgenome.org), with the default settings (Holm-Bonferroni correction).

#### Gene Property Analyses

To test for characteristic properties of Pufp3 cis targets, Puf3p trans effect proteins, and mitochondrial proteins, we compared our data against numerous publicly available data sets. Briefly, all proteins quantified in ∆*puf3* yeast (n = 3152, fermentation growth conditions, Y3K data set (Stefely et al., 2016a)) were assigned to one or more of the following categories where appropriate: Puf3p cis targets (n = 91, as defined in this report), Puf3p trans effect proteins (n = 49, as defined in this report), mitochondrial proteins (Jin et al., 2015) (n = 715), and all profiled proteins (n = 3152) in fermenting ∆*puf3* yeast. For each gene property analysis, each protein was assigned a qualitative or quantitative value as reported in a publicly available data set if a corresponding value was reported therein. Protein overexpression toxicity was assigned using data from (Gelperin et al., 2005). Enrichment in protein toxicity amongst cis, trans, and mitochondrial proteins was calculated relative to all proteins. Statistical significance of these enrichments was determined via a Fisher’s exact test. Protein abundance, in fermentation and respiration conditions, was assigned by taking the average log_2_ label free quantitation (LFQ) value from 36 replicates of WT yeast grown as part of the Y3K study (Stefely et al., 2016a). Data sets have been reported previously for cotranslational import of mitochondrial proteins (Williams et al., 2014), protein abundance changes across the diauxic shift (Stefely et al., 2016b), and average respiration deficiency response (RDR) protein abundance change (change in protein abundance in respiration deficient yeast compared to respiration competent yeast) (Stefely et al., 2016a). For these quantitative variable analyses, statistical significance was calculated using a Student’s *t*-test (two-tailed, homostatic).

#### Lipid Extractions

Frozen pellets of yeast (10^8^ cells) were thawed on ice and mixed with glass beads (0.5 mm diameter, 100 µL). CHCl_3_/MeOH (2:1, v/v, 4 °C) (900 µL) and CoQ_10_ (10 µL, 10 µM, 0.1 nmol) were added and vortexed (2 × 30 s). HCl (1 M, 200 µL, 4 °C) was added and vortexed (2 × 30 s). The samples were centrifuged (5,000 *g*, 2 min, 4 °C) to complete phase separation. 555 µL of the organic phase was transferred to a clean tube and dried under Ar_(g)_. The organic residue was reconstituted in ACN/IPA/H_2_O (65:30:5, v/v/v) (100 µL) for LC-MS analysis.

#### LC-MS Lipid Analysis

LC-MS analysis was performed on an Acquity CSH C18 column held at 50 °C (100 mm × 2.1 mm × 1.7 µm particle size; Waters) using a Vanquish Binary Pump (400 µL/min flow rate; Thermo Scientific). Mobile phase A consisted of 10 mM ammonium acetate in ACN/H_2_O (70:30, v/v) containing 250 µL/L acetic acid. Mobile phase B consisted of 10 mM ammonium acetate in IPA/ACN (90:10, v/v) with the same additives. Mobile phase B was held at 40% for 6.0 min and then increased to 60% over 3.0. Mobile phase B was further increased to 85% over 0.25 min and then to 99% for over 1.25 min. The column was then reequilibrated for 3.5 min before the next injection. Ten microliters of sample were injected by a Vanquish Split Sampler HT autosampler (Thermo Scientific). The LC system was coupled to a Q Exactive mass spectrometer by a HESI II heated ESI source kept at 325 °C (Thermo Scientific). The inlet capillary was kept at 350 °C, sheath gas was set to 25 units, and auxiliary gas to 10 units, and the spray voltage was set to 3,000 V. The MS was operated in positive and negative parallel reaction monitoring (PRM) mode acquiring scheduled, targeted PRM scans to quantify key CoQ intermediates. Phospholipids were quantified and identified using a negative dd-Top2 scanning mode.

*LC-MS Lipid Data Analysis.* CoQ intermediate data were processed using TraceFinder 4.0 (Thermo Fisher Scientific). Discovery lipidomic data were processed using an in-house software pipeline and Compound Discoverer 2.0 (Thermo Fisher Scientific).

#### Fluorescence Microscopy

Yeast (1×10^8^ cells) transformed with various FLAG tagged constructs were removed from cultures by pipetting and immediately fixed with formaldehyde (4% final concentration, gentle agitation on a nutator, 1 h, ~23 °C). The fixed cells were harvested by centrifugation (1000 *g*, 2 min, ~23 °C), washed three times with 0.1 M potassium phosphate pH 6.5 and once with K-Sorb buffer (5 mL, 1.2 M sorbitol, 0.1 M KP_i_, pH 6.5), and re-suspended in K-Sorb buffer (1 mL). An aliquot of the cells (0.5 mL) was mixed with K-Sorb-BME (0.5 mL, K-Sorb with 140 mM BME) and incubated (~5 min, ~23 °C). Zymolase 100 T was added to 1 mg/mL final concentration and incubated (20 min, 30 °C). The resultant spheroplasts were harvested by centrifugation (1000 *g*, 2 min, ~23 °C), washed once with K-Sorb buffer (1.4 mL), and re-suspended in K-Sorb buffer (0.5 mL). A portion of the cells (0.25 mL) was pipetted onto a poly-D-lysine coated microscope coverslip and allowed to settle onto the slides (20 min, ~23 °C). To permeabilize the cells, the supernatant was aspirated from the coverslips, and MeOH (2 mL, −20 °C) was added immediately and incubated (6 min, on ice). The MeOH was aspirated and immediately replaced with acetone (2 mL, −20 °C) and incubated (30 s, on ice). The acetone was aspirated, and the slides were allowed to air-dry (~2 min). DNA was stained with Hoechst 33342 dye (1 μg/mL in PBS, 2 mL, 5 min incubation, protected from light), and the cells were immediately wash with PBS. The samples were blocked with BSA (“BSA-PBS” [1% BSA in PBS], 2 mL, 30 min incubation at ~23 °C), and incubated with primary antibodies (1 mg/mL stock anti-FLAG primary Ab [Sigma F1804] at a 1:2000 dilution in PBS-BSA; anti-Cit1p antibody [Biomatik Anti-SA160118(Ser)] at a 1:500 dilution in PBS-BSA; 1 mL, 12 h, 4 °C). The anti-Cit1p antibody used in this experiment was generated against the peptide CRPKSFSTEKYKELVKKIESKN (Biomatik, AB001455, Anti-SA160118(Ser), lot # A160414-SF001455, peptide 506543, rabbit RB7668, 0.52 mg/mL). The samples were washed 5 times with PBS-BSA (2 mL, ~23 °C) and incubated with secondary antibodies diluted in PBS-BSA [1 μg/mL working concentration for each: Goat anti-Mouse IgG (H+L) Secondary Antibody, Alexa Fluor 594 conjugate (Thermo A-11005) and Goat anti-Rabbit IgG (H+L) Secondary Antibody, Alexa Fluor 488 (Thermo Cat# A-11008)] (1 mL, 2 h, ~23 °C, in the dark). The samples were washed 5 times with PBS-BSA (2 mL, ~23 °C) and twice with PBS (2 mL, ~23 °C). The last wash was aspirated and the slides were allowed to air dry briefly in the dark. The coverslips were mounted onto slides with 50% glycerol in PBS (8 μL). Fluorescence microscopy was performed on a Keyence BZ-9000 microscope using 100X oil immersion optics at room temperature. Line scan analysis was performed with ImageJ.

### QUANTIFICATION AND STATISTICAL ANALYSIS

#### Overview of Statistical Analyses

For each reported *P*-value, the statistical test used is reported in the legend for the corresponding figure panel. The majority of *P*-values in this report were calculated using an unpaired, two-tailed, Student’s *t*-test. In a few select instances, as noted, *P*-values for hypergeometric tests and Spearman correlation coefficients were calculated using the *R* software suite. Also as noted above, a Fischer’s exact test was used for a few select qualitative gene set analyses. For yeast experiments, all instances where n replicates are reported had n biological replicates. As detailed above, for the gene and protein set analyses, n indicates the number of genes or proteins in each set: Puf3p cis targets (n = 91), Puf3p trans effect proteins (n = 49), mitochondrial proteins (n = 715), and all profiled proteins (n = 3152) in fermenting ∆*puf3* yeast.

